# Animal-free recombinant nanobody rescues HCN4 channel deficit in sinus node dysfunction

**DOI:** 10.1101/2025.03.20.644287

**Authors:** Atiyeh Sadat Sharifzadeh, Roberta Castelli, Alessandro Porro, Pietro Mesirca, Romain Perrier, Ana M Gómez, Hugo Benoit, Albano C. Meli, Luca M G Palloni, Dario DiFrancesco, Matteo E. Mangoni, Gerhard Thiel, Andrea Saponaro, Anna Moroni

**Author notes:** Equally contributed (Atiyeh Sadat Sharifzadeh, Roberta Castelli).

## Abstract

Hyperpolarization-activated cyclic nucleotide-gated channels (HCN1-4) control cardiac and neuronal firing and their dysfunction leads to cardiac arrythmias (HCN4), epilepsy (HCN1) and chronic pain (HCN2). Prompted by the urgent need for HCN subtype-specific treatments, we screened a recombinant nanobody library in search of HCN4-specific binders. Here we show that nanobody 5 (NB5) binds to the extracellular side of HCN4 with high specificity and nanomolar affinity and activates the channel by a non-canonical electromechanical coupling path. In *ex vivo* and *in vitro* experiments, NB5 acts as an agonist of the pacemaker current I_f_, increasing the firing rate of rabbit cardiac pacemaker myocytes and of human derived cardiomyocytes. Notably, NB5 rescued the effects of a LOF HCN4 mutation causing sinus node dysfunction in a patient. Our work illustrates that animal-free recombinant nanobodies have strong potential as next generation modulators for clinical application in symptomatic bradycardia.

## Introduction

Hyperpolarization-activated and cyclic nucleotide-gated (HCN) channels are encoded by four human genes (*hcn1-4*) which are widely expressed, in a cell type specific manner, in the heart and in the nervous system. Their activation by membrane hyperpolarization and cyclic nucleotides underlies the so-called pacemaker current I_*f/h*_ which regulates the rhythmicity of cardiac (I_*f*_) and neuronal (I_*h*_) excitation^1^. Since HCNs are involved in cardiac and neuronal pacemaking, channel dysfunctions lead to severe medical conditions. Depending on the subtype composition of the functional channels and their organ specific expressions, HCN-related pathologies range from disease of heart automaticity (sinus node dysfunction, HCN4) to neurological disorders, including the life-threatening Early Infantile Epileptic Encephalopathy (EIEE, HCN1). Furthermore, in the peripheral nervous system, HCN2 plays a major role in chronic pain^2–4^. Despite their pathophysiological importance, to date, only one clinically available drug targets HCN channels, the open pore blocker Ivabradine, which is indicated for *angina pectoris*^5^ and heart failure. Ivabradine is poorly subtype specific thus its use for the treatment of HCN1 and HCN2-based diseases unavoidably leads to the slowing of the heartbeat^6^. The recent cryo-EM structure of Ivabradine bound to HCN4 further revealed that the residues forming the main contacts to the drug inside the open pore are highly conserved in all HCNs^7^, suggesting that subtype specific drugs may likely emerge from alternative approaches rather than from chemical modifications of Ivabradine itself.

A promising alternative approach for developing subtype specific HCN drugs is provided by nanobodies (NBs). These small (15 kDa) and rather stable peptides, bind with high affinity to their epitopes and are currently emerging as a promising alternative to small chemical molecules in therapy^8,9^. Originally isolated from camelids, NBs can also be produced synthetically. By screening a synthetic nanobody library^10^ against the purified HCN4 protein, we have isolated an HCN4-specific nanobody, NB5, which binds to the extracellular side of the channel with nanomolar affinity. NB5 is a “functional” nanobody as it activates the channel by inducing a depolarizing shift of its voltage dependency by about 10 mV. Proof of concept experiments confirm that NB5 increases pacemaker activity in native rabbit sino-atrial node (SAN) myocytes and cardiomyocytes differentiated from human pluripotent stem cells. Finally, NB5 fully reverses the hyperpolarized phenotype of a mutant HCN4 channel found in a patient with a genetic cardiac disorder.

## Results

### Nanobodies isolation from the library

We screened a yeast-display nanobody library^10^ against rabbit HCN4 (HCN4), detergent-purified from HEK293F cells in two conformations, with the pore open (HCN4_open_) or closed (HCN4_closed_)^7,11^. The protein assembles as tetramer and each monomer includes all known functional domains of HCN channels, N-terminal HCN domain (HCND), voltage sensor (VSD, TM1-TM4), pore (PD, TM5-6), C-linker and Cyclic Nucleotide Binding Domain (CNBD) with downstream helices D and E^12^, but lacks residues 783 - 1064 in the C terminus and has an eGFP fused at the N terminus (GFP-HCN4). Yeast cells displaying positive nanobodies - nanobodies binding the HCN4 protein - were isolated by iterative cycles of magnetic activated cell sorting (MACS), followed by fluorescence activated cell sorting (FACS) (Figure S1a). In MACS, positive cells were retrieved using anti-GFP antibody attached to magnetic beads; in FACS, positive cells were sorted by following the fluorescent signal of GFP- HCN4. The screening procedure included several enrichment cycles, with HCN4_OPEN_ as an antigen, and depletion cycles, with HCN4_CLOSED_ as an antigen (Figure S1b). The latter were added to enrich the library of open-state binders, in search of potential pore blockers. FACS analysis was conducted in parallel to monitor the progressive enrichment of the library in HCN4 binders, from the initial 5% to the final 37% (Figure S1b). At the end of the screening, the recovered yeast cells were plated, and 100 colonies were randomly picked and sequenced. Their sequencing returned ten independent clones (Figure S2a,b) that were expressed in *E*.*coli* and purified as His-tag proteins.

### Functional test of nanobodies on HCN channels

We next evaluated in patch clamp experiments whether the purified NBs had any effect on channel function. Wild type rabbit HCN4 (rbHCN4) channels were expressed in HEK293T cells, and the NBs (20 µM) were added to the extracellular solution 2-3 minutes before patching. In the second round of screening, the NBs that did not display any effect from the extracellular side (p values >0.05) were added to the pipette solution and further tested from the intracellular side. The results, summarized in Table S1, show that six NBs modified channel function acting from the extracellular side, while 4 did not have any effect either from the extracellular or intracellular side, or could not be tested because they were unstable in solution. All six active NBs induced a right shift in the activation curve, ranging from 3 to 13 mV, with NB5 and NB6 being the most soluble and stable proteins.

To check for subtype-specificity, the six active NBs were also tested on hHCN1, mHCN2 and hERG, an HCN related member of the Kv superfamily^13^. Two NBs, namely NB3 and NB5, were inactive on hHCN1. NB5 was also inactive on mHCN2 and hERG and neither changed the conductance nor the voltage-dependency of these two channels (Figure S3 and Table S2), confirming its specificity for rbHCN4. Furthermore, we found that NB6 was active on hHCN1 (ΔV_1/2_ = 8.4 ± 1.7 mV) but had no effect on mHCN2 and hERG (Figure S4 and Table S2). Finally, NB66 acted on all channels, indicating a non-specific mechanism of action or an artefactual result.

### In-depth characterization of NB5 effect on rbHCN4

NB5 increased the current at intermediate voltages, as shown by the increase at -105 mV (Figure 1a, black arrowhead), but not the maximal current at -150 mV (Figure 1b). This effect was accompanied by acceleration of the activation kinetics, while deactivation did not change (Figure 1a). This behavior is indicative of a right-shift in the voltage-dependent activation of the channel, shown in Figure 1c. Fitting data to a Boltzmann equation (solid lines) yielded the half-activation voltage values (V_1/2_) reported in Figure 1d. Mean values ± SEM are as follow: ctrl = -104.8 ± 0.8 mV, +NB5 = -95.7 ± 0.9 mV. The NB5-induced shift (ΔV_1/2_) ± SEM is 9.1 ± 1.2 mV (Table S1). Fitting the dose-response data of NB5 (Figure 1e) to a Hill function (solid line) yielded half maximal effective concentration (K_1/2_) value of 40 nM. The Hill coefficient (n_H_) value is 0.65. The effect of NB5 persisted during washout for at least 70 minutes (Figure S5 and Table S3), confirming the high affinity of NB5 for the channel.

**Figure 1.**
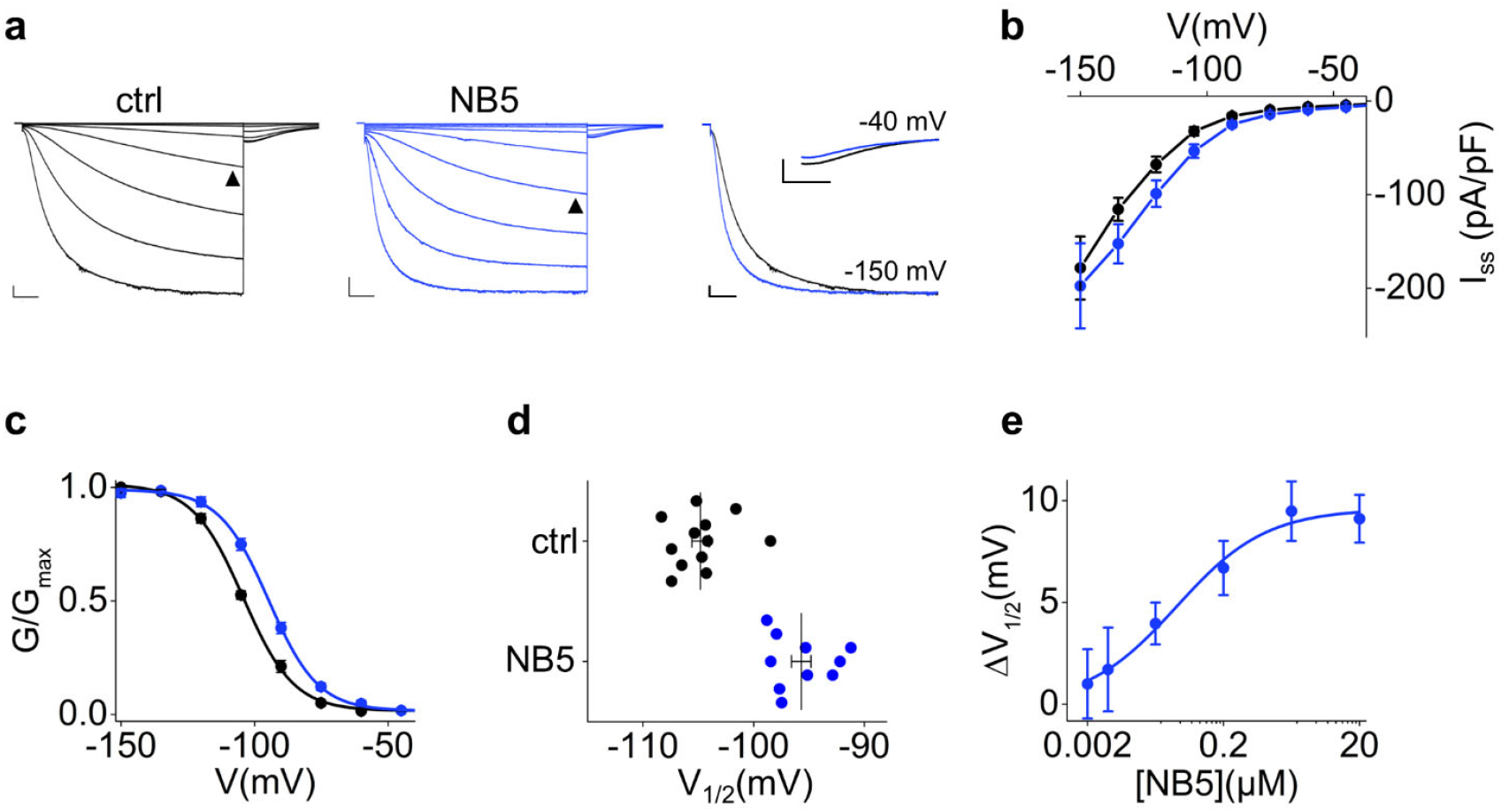
Functional effect of NB5 on rbHCN4. **a**. Representative whole-cell currents of wt rbHCN4 expressed in HEK293T cells and recorded without (left) or with (middle) 20 µM NB5 in the extracellular solution. Traces shown are from -30 mV to -150 mV (see Material and Methods for voltage step protocol). Black arrowheads indicate current at -105 mV. The right panel shows superposition of traces recorded from a cell without (black trace) and a cell with NB5 (blue trace) at -150 mV and tail currents collected at -40 mV. Scale bars: 250 pA and 500 ms. **b**. Mean I/V relationships of steady state current density (I_ss_, pA/pF) obtained from cells with (blue) or without (black) 20 µM NB5 in the extracellular solution. Data are mean of n=12 cells (ctrl) and n=10 (NB5) ± SEM. **c**. Activation curves obtained from cells in b. Data fit to the Boltzmann equation (see Material and Methods) are plotted as solid lines. Data points are mean ± SEM. Individual V_1/2_ values are shown in d. **d**. Half-activation voltages (V_1/2_) of individual cells plotted in c. Mean V_1/2_ ± SEM for ctrl = -104.8 ± 0.8 mV, +NB5 = -95.7 ± 0.9 mV. The shift in half activation voltage (ΔV_1/2_) ± SEM = 9.1 ± 1.2 mV. **e**. ΔV_1/2_ (mV), measured as in d, in response to a range of NB5 concentrations (µM). Data fit to the Hill equation (see Material and Methods) (solid line) yielded a half maximal effective concentration (K_1/2_) value of 40 nM and a Hill coefficient (n_H_) value of 0.65. Data are mean ± SEM. Each data point is an average of n ≥ 3 cells. V_1/2_, ΔV_1/2_, inverse slope factors (k) and number of cells (n) for each experiment shown are reported in Table S1 along with the details on statistical analysis.

### Identification of NB5 binding site

The list of potential binding sites of NB5 on HCN4 channels includes the extracellular side of TM1-6 helices and the short loops connecting them (S1-S2, S3-S4 and S5-PH, PH-S6). We narrowed them down at first, based on the subtype-specificity of NB5. Alignment of rbHCN4 with hHCN1 and mHCN2 shows HCN4-specific residues in five positions, two in the VSD and three in the PD (Figure S6).

However, the precise identification of the residues involved in NB5 binding was greatly facilitated by the finding that NB5 is not only subtype-but also ortholog-specific. Indeed, NB5 activates human (hHCN4) but not mouse HCN4 (mHCN4) (Figure 2a; Table S3). Thus, alignment of the ortholog sequences identified in the S5-P-helix loop two residues, D448 and N456, that are conserved in rabbit and human, but not in mouse HCN4 (Figure 2b). Mapping of the two residues on the structures of rbHCN4 (7NMN, 7NP4, open and closed pore, respectively) and of hHCN4 (6GYO, closed pore) shows that they are in the pore turret and are freely accessible from the extracellular solution (Figure 2c). We validated the binding site by introducing the mouse residues (D448H and N456G), alone or in combination, in rabbit HCN4. Both mutations did not alter the voltage dependency of the channel, as shown from the V_1/2_ of the mutants that is the same as the wt but greatly reduced (D448H), or fully abolished (D448/N456G), the response to NB5 (Figure 2d, Table S3). From the dose-response curves, we calculated a K_1/2_ of 400 nM for D448H, a 10-fold drop in NB5 affinity compared to the wt, while the double mutant D448H/N456G was virtually NB5-insensitive (Figure 2e, Table S3).

**Figure 2.**
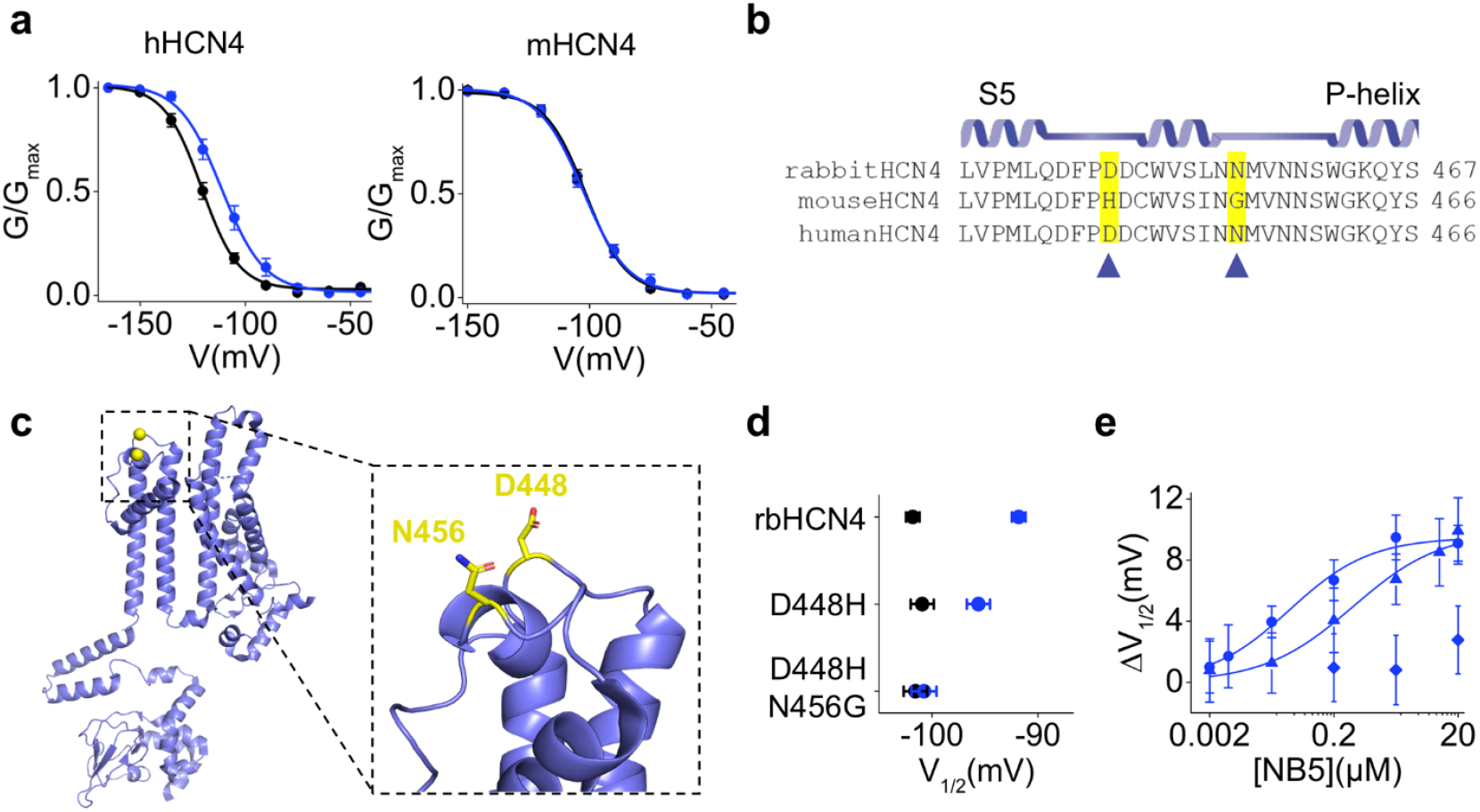
Identification of NB5 binding site on rbHCN4. **a**. Activation curves obtained from hHCN4 wt (left) and mHCN4 (right) currents in control solution (black circles) or in the presence of 2 µM NB5 in the extracellular solution (blue circles). Data fit to the Boltzmann equation (see Material and Methods) are plotted as solid lines. Data points are mean ± SEM. **b**. Secondary structure elements and multiple sequence alignment of the S5-Pore-helix region of rabbit HCN4 (Gene ID: 100009452), mouse HCN4 (Gene ID: 330953) and human HCN4 (Gene ID: 10021). Residues conserved in both rbHCN4 and hHCN4 are highlighted in yellow and are indicated by arrowheads. **c**. Ribbon representation of rabbit HCN4 monomeric structure in the open pore configuration (PDB: 7NP3^11^) with residues highlighted in panel a (D448 and N456) shown as yellow spheres. Boxed: blow-up showing D448 and N456 side chains. Numbering refers to rbHCN4 sequence. **d**. Mean half-activation voltages (V_1/2_) of control (black circles) and 2 µM NB5-treated (blue circles) cells expressing rbHCN4 wt, rbHCN4 D448H or rbHCN4 D448H_N456G channels. Data shown are mean ± SEM. NB5-induced shift in half activation voltage (ΔV_1/2_) ± SEM = 9.1 ± 1.2 mV (on wt), 6.7 ± 1.6 mV (on D448H) and 0.8 ± 2.3 mV (on D448H_N456G). **e**. ΔV_1/2_ (mV) plotted as a function of NB5 concentration (µM) in rbHCN4 wt (blue circles), rbHCN4 D448H (blue triangles) and rbHCN4 D448H_N456G (blue diamonds). Data fit to the Hill equation (see Material and Methods) (solid blue line) yielded a half maximal effective concentration (K_1/2_) value of 40 nM (wt) and 400 nM (D448H) and a Hill coefficient (n_H_) value of 0.65 (wt) and 0.61 (D448H). Data points for D448H_N456 construct could not be fitted with a curve. Data are mean ± SEM. Each data point is an average of n ≥ 3 experiments. V_1/2_, ΔV_1/2_, inverse slope factors (k) and number of cells (n) for each experiment shown are reported in Supplementary Table S3 along with the details on statistical analysis.

### NB5 allosteric pathway is independent from that of cAMP

The effect of NB5 resembles at first glance that of cAMP, the endogenous activator of HCN channels known to shift the channel activation curve to the right^14^. However, it is well known that cAMP acts allosterically on the VSD from the intracellular side^11,15^, while NB5 binds the extracellular turret, a so far unexplored domain in terms of allosteric effects on the VSD.

To understand if the two mechanisms are independent, we checked whether the effects of NB5 and cAMP are additive or synergic. Figure 3a shows rbHCN4 current traces in control solution (ctrl) and in the presence of saturating concentrations of NB5 (2 µM), cAMP (30 µM) and both. Analysis of the activation curves shows that the effects of the two ligands on the channel voltage dependency are perfectly additive (Figure 3b,c). The mean shift ΔV_1/2_ was 8.5 ± 2.1 mV with NB5, 18.0 ± 2.3 mV with cAMP and 28.5 ± 2.2 mV with both ligands present (Figure 3c and Table S4). Plotting time constants of activation and deactivation (τ) shows that here, too, cAMP is similarly active on both control and NB5-bound channels (Figure 3d and Table S5).

**Figure 3.**
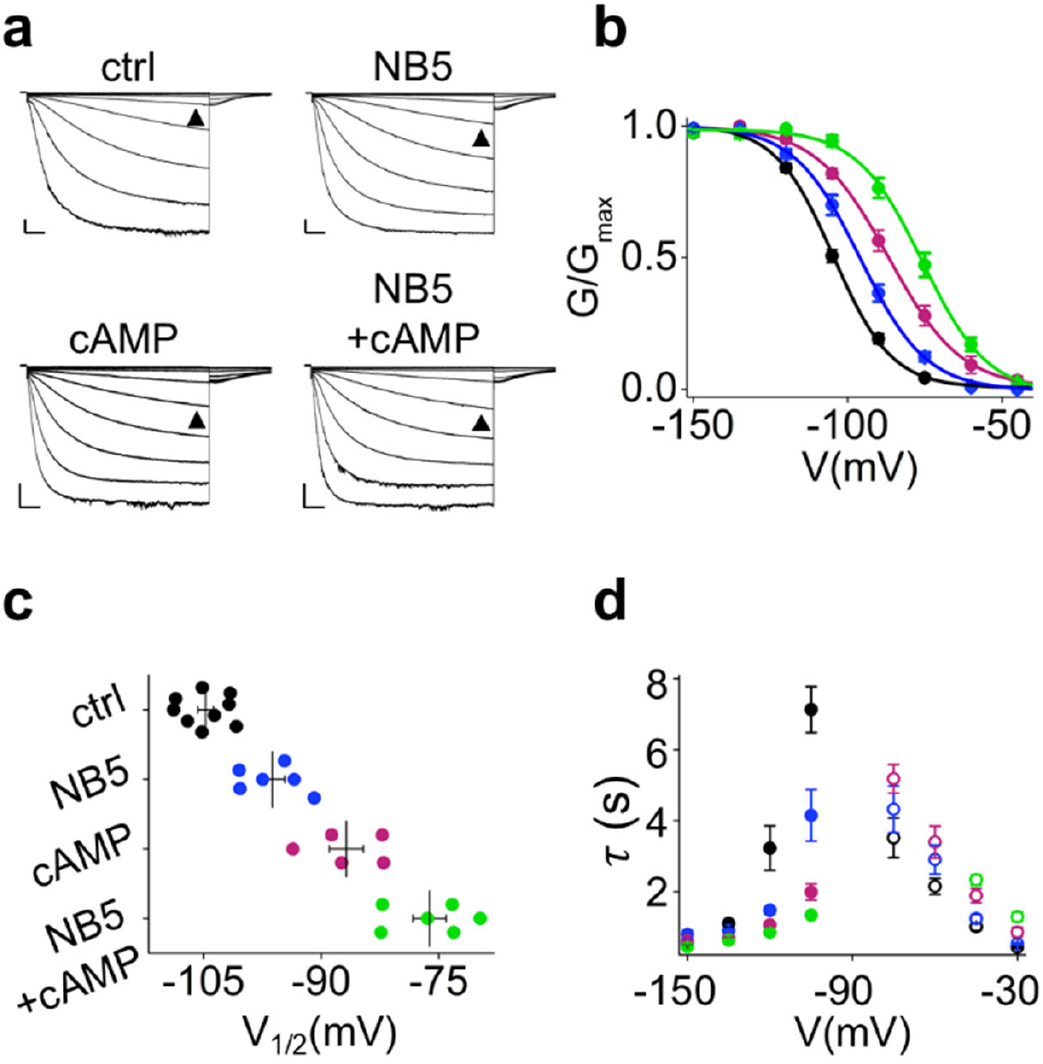
Effect of NB5 in the presence of cAMP. **a**. Representative whole-cell currents of rbHCN4 wt recorded in control solution (top left), in presence of 2 µM NB5 in the extracellular solution (top right), 30 µM cAMP in the pipette solution (bottom left) or both (bottom right). Traces shown are from -30 mV to -150 mV (see Material and Methods for voltage step protocol). Black arrowheads indicate current at -90 mV. Scale bars: 250 pA and 500 ms. **b**. Activation curves obtained from rbHCN4 wt in control solution (black circles) or in the presence of 2 µM NB5 in the extracellular solution (blue circles), of 30 µM cAMP in the pipette solution (pink circles) or both (green circles). Data fit to the Boltzmann equation (see Material and Methods) are plotted as solid lines. Data points are mean ± SEM. Individual V_1/2_ values are shown in c. **c**. Half-activation voltages (V_1/2_) of control (black circles, n=9), NB5 (blue circles, n=6), cAMP (pink circles, n=5) and NB5 + cAMP (green circles, n=6) -treated cells expressing rbHCN4 wt. Mean V_1/2_ ± SEM (in mV) for ctrl = -104.7 ± 1.0, +NB5 = -96.2 ± 1.6, +cAMP = -86.8 ± 2.2, +NB5 +cAMP = -76.2 ± 2.1. **d**. Mean activation (solid circles) and deactivation (empty circles) time constants (τ) of rbHCN4 wt in control solution (black), or in the presence of 2 µM NB5 (blue), 30 µM cAMP (pink) or both (green). Time constants were calculated, at the indicated voltages, by fitting a single exponential function (see Material and Methods) to current traces of panel a (activation) and to current traces obtained with a deactivating protocol (deactivation, see Material and Methods). Data points are mean ± SEM. Each data point is an average of n ≥ 3 experiments. V_1/2_, ΔV_1/2_, inverse slope factors (k), kinetics (τ_act_, τ_deact_) and number of cells (n) are reported in Supplementary Tables S4 and S5 along with the details on statistical analysis.

This finding shows that voltage-dependent modulation of NB5 is independent from cAMP mechanism of action, uncovering a non-canonical electromechanical coupling between the VSD and the pore turret, so far unexplored in HCN channels.

### State-dependent binding

Even though it seems likely that NB5 binds to the two residues in the turret, we cannot exclude that its binding site includes the VSD. Thus, we set up an experiment to test if NB5 binding is state-dependent. In other words, we know already that NB5 binds to the closed channel, but we wanted to evaluate if the open channel conformation further promotes its binding affinity.

To study the state dependence of NB5 action, we evaluated if the effect of NB5 on activation kinetics (τ_act_) increases over time, using a repetitive activation protocol (1/30Hz)^16^. NB5 was provided at a saturating concentration on the closed channel, while activation was performed at the half activation voltage (-90 mV). If the affinity of NB5 further increases in the open state, we would expect a time-dependent change in τ_act_ as more open channels become available. On the contrary, if NB5 does not discriminate between the closed and open conformation of the VSD, its effect would be completed in the first sweep. Analysis of the activation kinetics before and after NB5 treatment shows that the effect of the NB does not significantly change after the first sweep (Figure S7a,b) providing no evidence for state dependent binding. This apparently excludes the VSD leaving the pore domain as the only binding site.

### Effect of NB5 on cardiac sinoatrial node cells

To unfold its therapeutic potential, we tested whether NB5 is active on the native current (I_f_) of SAN myocytes. To this end, we conducted preliminary experiments to investigate if NB5 recognizes heteromeric channels composed of HCN4 and HCN1, that are likely to exist in SAN myocytes^17^. Heteromeric channels were formed in HEK293T cells by co-transfection of rbHCN4 and hHCN1 plasmids (Figure 4) and showed a V_1/2_ value of 84.8 ± 1.6 mV, intermediate between that of HCN4 and HCN1, and activation kinetics ten-times faster than those of HCN4 (τ_act_ HCN1/4 = 0.6 s vs HCN4 = 7.1 s at -105 mV, Table S5). Addition of NB5 increased the current at non saturating potentials (Figure 4a,b) by shifting the activation curve to the right (ΔV_1/2_ = 12.7 ± 2.5 mV) (Figure 4c,d) and accelerated the activation kinetics of the heteromeric channels (Figure 4e). Interestingly, the effect of NB5 on the heterotetrametric channel resembles that on HCN4, indicating that the 1:1 stoichiometry (NB5: monomer) is not a strict requirement for its action.

**Figure 4.**
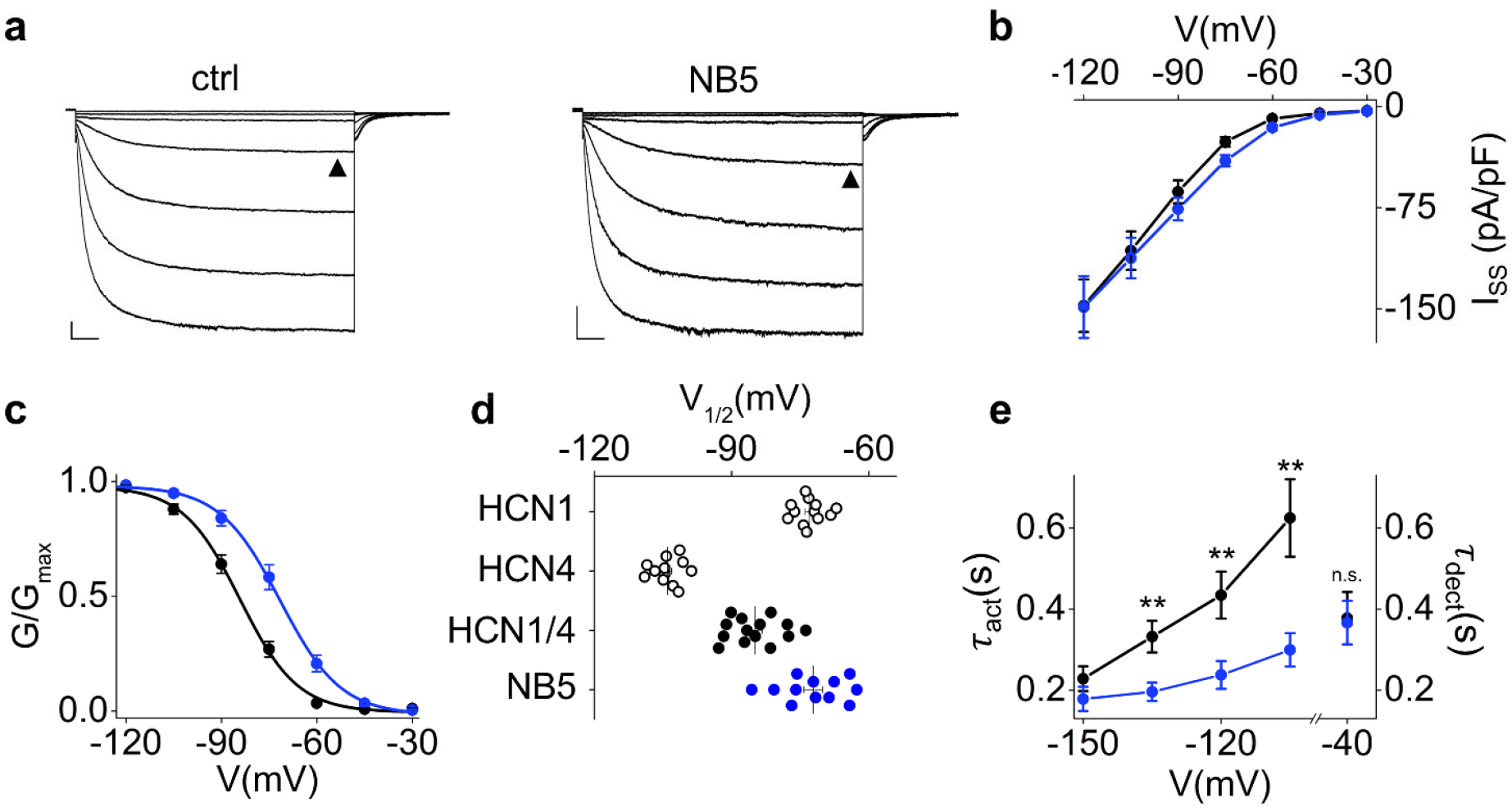
Effect of NB5 on hHCN1 / rbHCN4 heterotetramers. **a**. Representative whole-cell currents of hHCN1 wt / rbHCN4 wt (co-transfected, see Material and Methods) recorded in control solution (left) or in the presence of 2 µM NB5 in the extracellular solution (right). Traces shown are from -30 mV to -120 mV (see Material and Methods for voltage step protocol). Black arrowheads indicate current at -75 mV. Scale bars: 250 pA and 500 ms. **b**. I/V relationships of hHCN1 wt / rbHCN4 wt in control solution (black) or in the presence of 2 µM NB5 in the extracellular solution (blue). Data are mean ± SEM. Current density values (in pA/pF), indicated as I_SS_ (current at steady state), recorded at -120 mV, HCN1/4 = -147.7 ± 19.4 (n = 10), +NB5 = -148.8 ± 22.9 (n = 10) are not statistically different (^§^p=0.4, Student’s T-test). **c**. Activation curves obtained from hHCN1 wt / rbHCN4 wt currents in control solution (black circles) or in the presence of 2 µM NB5 in the extracellular solution (blue circles). Data fit to the Boltzmann equation (see Material and Methods) are plotted as solid lines. Data points are mean ± SEM. Individual V_1/2_ values are shown in d. **d**. Half-activation voltages (V_1/2_) of control (black circles, n=14) and NB5-treated (blue circles, n=12) cells expressing hHCN1 wt / rbHCN4 wt. Mean V_1/2_ ± SEM for HCN1/4 = -84.8 ± 1.6 mV, +NB5 = -72.2 ± 2.0 mV. Shift in half activation voltage (ΔV_1/2_) ± SEM = 12.7 ± 2.5 mV. Half-activation voltages (V_1/2_) of cells expressing either hHCN1 (n=12, V_1/2_ = -73.0 ± 0.9 mV) or rbHCN4 (n=12, V_1/2_ = -104.0 ± 0.9 mV) in control solution are represented as empty black circles. **e**. Mean activation and deactivation time constants (τ_act_, τ_deact_) of hHCN1 wt / rbHCN4 wt in control solution (black circles) or in the presence of 2 µM NB5 (blue circles). Time constants were calculated, at the indicated voltages, by fitting a single exponential function (see Material and Methods) to current traces of panel a. Data points are mean ± SEM. Each data point is an average of n ≥ 3 experiments. Statistical analysis performed with Student’s two-tailed unpaired T-test compared to control condition (**p < 0.01). V_1/2_, ΔV_1/2_, inverse slope factors (k), kinetics (τ_act_, τ_deact_) and number of cells (n) are reported in Supplementary Tables S3 and S5 along with the details on statistical analysis.

NB5 was then tested on myocytes isolated from rabbit and mouse hearts, the latter was used as a negative control. Figure 5a-c shows that NB5 did not affect I_*f*_ recorded from mouse SAN (n = 4), while it affected the current recorded from rabbit SAN (Figure 5d-f; Table S3) shifting V_1/2_ in average by +11.6 ± 5.1 mV (n = 7). Most importantly, the action potential firing rate of rabbit SAN myocytes was increased by NB5 (Figure 5g) in 6 out of 9 cells tested (Figure 5h). On average, spontaneous beating was heightened by 16 bpm, corresponding to 30% increase in the relative beating rate. These results confirm that NB5 activates the native f-channel of rabbit pacemaker myocytes, suggesting that NB5 may harbor therapeutic potential to improve pacemaker activity in human sinus node dysfunction. To test this hypothesis, we investigated the potency of NB5 in activating HCN4 in spontaneously beating cardiomyocytes, differentiated from human pluripotent stem cells (hiPSC-CMs). These myocytes are known to express HCN4, which facilitates their spontaneous beating rate^18^. We obtained hiPSC-CMs differentiated into a 2D cardiac sheet. Using video-edge capture, we recorded the contractility properties of the cardiac sheet’s contractions before and after the application of NB5, allowing for a paired comparison. Our results indicated that NB5 incubation significantly increased sheet beating rate, as evidenced by reduced systolic and diastolic periods (Figures 5i, S8a,b). In line with the hypothesis of selective effect on HCN4, perfusion of NB5 left contraction amplitude or contraction heterogeneity unaffected, thereby eliminating the presence of arrhythmic events induced by NB5 application (Figure S8c,d).

**Figure 5.**
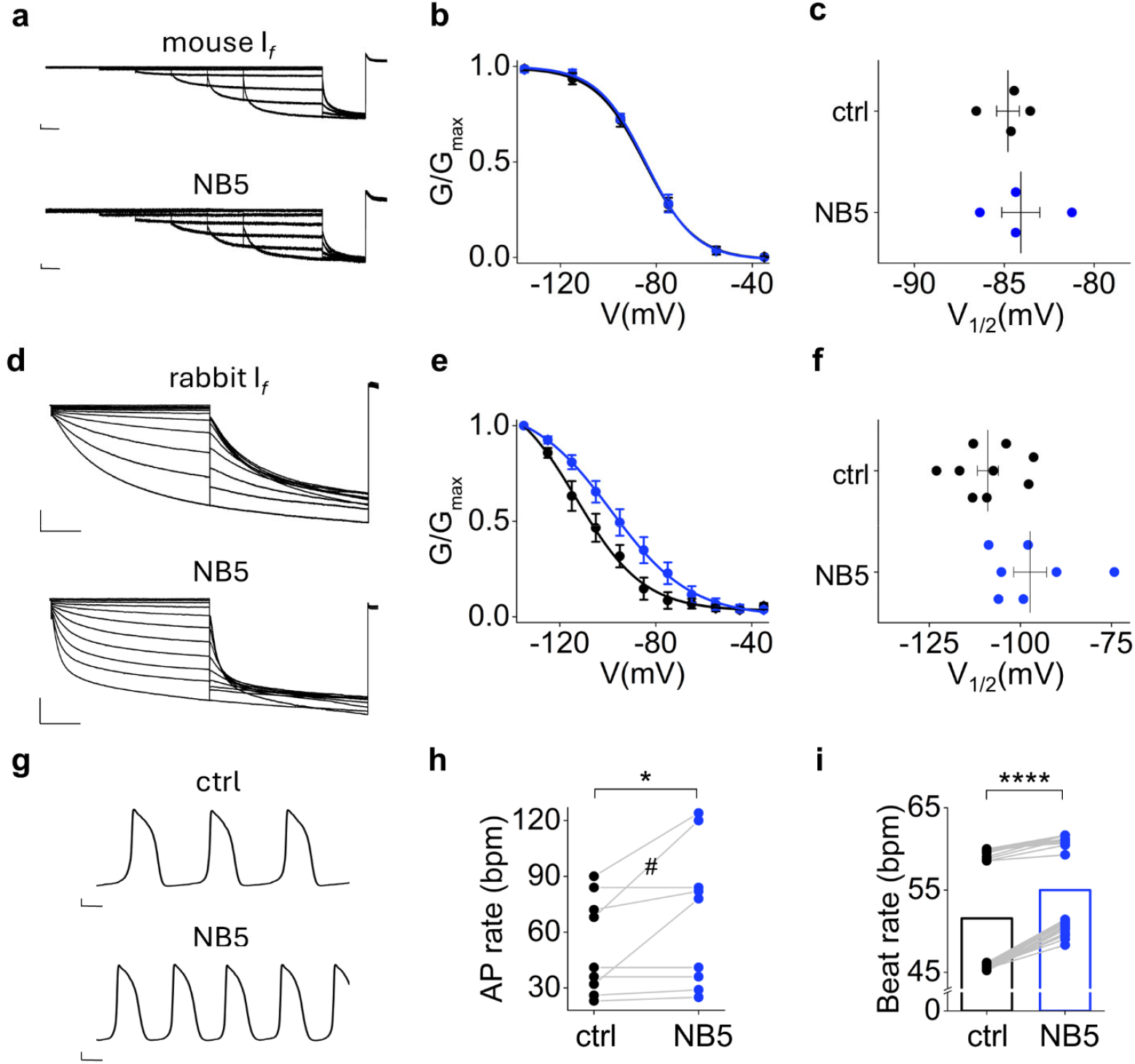
Effect of NB5 on voltage-dependent activation of I_*f*_ and spontaneous rate in cardiac sinoatrial node (SAN) cells. **a**. Representative whole-cell I_*f*_ currents recorded from mouse SAN cells in control solution (left) or pre-incubated with 4 µM NB5 in the extracellular solution for 5 minutes (right). Traces shown are from -35 mV to -135 mV (see Material and Methods for voltage step protocol). Scale bars: 250 pA and 500 ms. **b**. I_*f*_ activation curves obtained from mouse SAN currents in control solution (black circles) or pre-incubated with 4 µM NB5 in the extracellular solution (blue circles). Data fit to the Boltzmann equation (see Material and Methods) are plotted as solid lines. Data points are mean ± SEM. Individual V_1/2_ values are shown in c. **c**. Half-activation voltages (V_1/2_) of control (black circles, n=4) and NB5-treated (blue circles, n=4) mouse SAN cells. Mean V_1/2_ ± SEM for ctrl = -84.8 ± 0.6 mV, +NB5 = -84.1 ± 1.1 mV. The shift in half activation voltage (ΔV_1/2_) ± SEM = 0.7 ± 1.2 mV. **d**. Representative whole-cell I_*f*_ currents recorded from rabbit SAN cells in control solution (left) or pre-incubated with 2 µM NB5 in the extracellular solution for 5 minutes (right). Traces shown are from -35 mV to -135 mV (see Material and Methods for voltage step protocol). Scale bars: 250 pA and 500 ms. **e**. I_*f*_ activation curves obtained from rabbit SAN currents in control solution (black circles) or pre-incubated with 2 µM NB5 in the extracellular solution (blue circles). Data fit to the Boltzmann equation (see Material and Methods) are plotted as solid lines. Data points are mean ± SEM. Individual V_1/2_ values are shown in f. **f**. Half-activation voltages (V_1/2_) of control (black circles, n=9) and NB5-treated (blue circles, n=7) rabbit SAN cells. Mean V_1/2_ ± SEM for ctrl = -108.9 ± 2.9 mV, +NB5 = 97.3 ± 4.5 mV. The shift in half activation voltage (ΔV_1/2_) ± SEM = 11.6 ± 5.1 mV. **g**. Representative recording of single rabbit SAN cell spontaneous activity before (top) and after (bottom) superfusion of 2 µM NB5. Scale bars: 10 mV and 2.5 sec. **h**. Spontaneous action potential rate (bpm) of single rabbit SAN cells before (black circles) and after (blue circles) superfusion of 2 µM NB5. # symbol indicates the representative cell shown in panel g. Mean AP rate ± SEM for ctrl = 52.3 ± 8.7 bpm, +NB5 = 68.7 ± 12.7 bpm. Statistical analysis performed with Student’s paired t-test (p=0.05). V_1/2_, ΔV_1/2_, inverse slope factors (k) and number of cells (n) for experiments of panel a-f are reported in Supplementary Table S3 along with the details on statistical analysis. **i**. Before-after graph of the beat rate (bpm) of spontaneously beating cardiac sheets containing hiPSC-CMs before (ctrl) and after addition of 2 µM NB5 (NB5) at 37°C. Empty columns show mean values. Statistical analysis performed with Wilcoxon paired test (****p < 0.0001).

### NB5 rescues a mutation of a sinus node dysfunction patient

Given its role in the generation of spontaneous activity in pacemaker cells, it is not surprising that mutations of the *hcn4* gene are associated with sinus node dysfunction (SND), leading to symptomatic sinus bradycardia^19^ and more complex lethal arrhythmias^4,20^. Most of the mutations described so far are loss of function. Functional loss is caused either by a negative shift of the activation curve or by lower density of membrane expression of channels and consequent reduction of current density^20^.

To test the therapeutic potential of NB5, we selected mutation K530N, previously reported in a SND patient eventually implanted with an artificial pacemaker because of symptomatic bradyarrhythmia (35 bpm)^21^. Lysine 530 is found in the A’ helix of the C-linker and its mutation into asparagine induces, in the heterozygous condition of the patient, slower activation kinetics and a left shift in V_1/2_ of about 14 mV^21^. We inserted the mutation in rabbit HCN4 (K531N) and confirmed the phenotype for the heterozygote channels wt/K531N (Figure 6a-d; Table S3). Addition of 0.2 µM NB5 shifted the activation curve to the right and reversed the phenotype of the mutant, both in terms of V_1/2_ and kinetics (Figure 6a-c; Tables S3 and S5). Notably, the effect of NB5 could be adjusted to the exact shift, as it was dose dependent (Figure 6d; Table S3).

**Figure 6.**
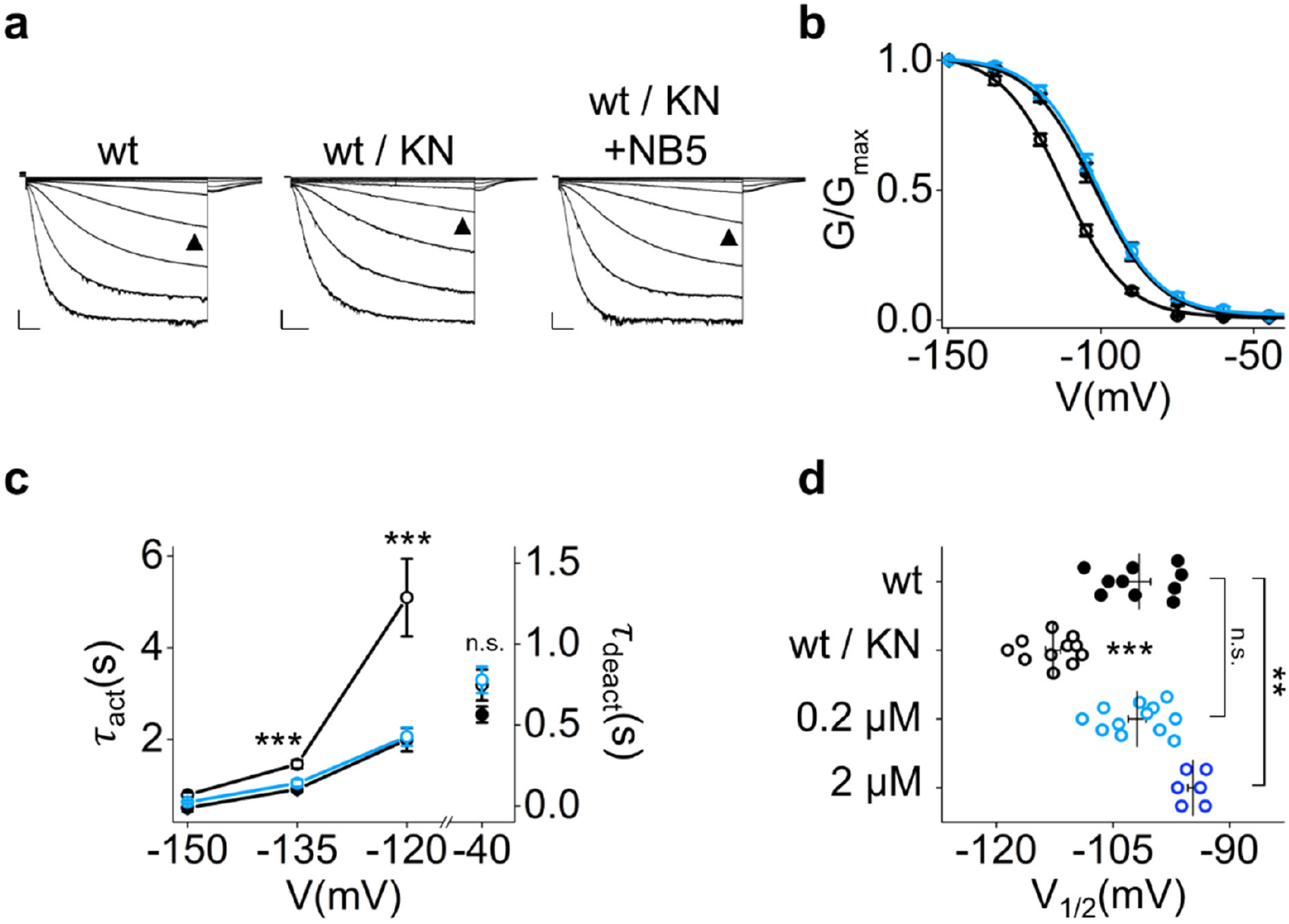
Effect of NB5 on HCN4 pathogenic variant K531N. **a**. Representative whole-cell currents of rbHCN4 wt (left) and wt / K531N (centre) recorded in control solution or in the presence of 0.2 µM NB5 in the extracellular solution (wt / K531N, right). Traces shown are from -30 mV to -150 mV (see Material and Methods for voltage step protocol). Black arrowheads indicate current at -105 mV. Scale bars: 250 pA and 500 ms. **b**. Activation curves obtained from rbHCN4 wt (black solid circles) and wt / K531N (black empty circles) currents in control solution or in the presence of 0.2 µM NB5 in the extracellular solution (wt / K531N, light blue empty circles). Data fit to the Boltzmann equation (see Material and Methods) are plotted as solid lines. Data points are mean ± SEM. Individual V_1/2_ values are shown in d. **c**. Mean activation and deactivation time constants (τ_act_, τ_deact_) of rbHCN4 wt (black solid circles) and wt / K531N (black empty circles) in control solution or in the presence of 0.2 µM NB5 (wt / K531N, light blue empty circles). Time constants were calculated, at the indicated voltages, by fitting a single exponential function (see Material and Methods) to current traces of panel a. Data points are mean ± SEM. Each data point is an average of n ≥ 3 experiments. Statistical analysis performed with Student’s two-tailed unpaired T-test compared to control condition (***p <0.001). **d**. Half-activation voltages (V_1/2_) of control wt (black solid circles, n=10), control wt / K531N (black empty circles, n=11) and wt / K531N cells treated with 0.2 µM NB5 (light blue empty circles, n=12) or 2 µM NB5 (dark blue empty circles, n=6). Mean V_1/2_ ± SEM for wt ctrl = -104.8 ± 0.8 mV, wt / K531N ctrl = -112.7 ± 1.0 mV, wt / K531N + 0.2 µM NB5 = -101.9 ± 1.1 mV, wt / K531N + 2 µM NB5 = -94.7 ± 0.7 mV. V_1/2_, ΔV_1/2_, inverse slope factors (k), kinetics (τ_act_, τ_deact_) and number of cells (n) for each experiment shown are reported in Supplementary Tables S3 and S5 along with the details on statistical analysis.

## Discussion

In this work we demonstrate that it is possible to develop subtype-specific nanobodies for HCN channels and that by acting from the extracellular side, they bear great potential for clinical applications. This finding promises to revolutionize the therapy of several HCN-related human diseases, many being currently without treatment.

Among ten NB candidates functionally characterized here by patch clamp, NB5 is the most suitable for further drug development because it is strictly HCN4-selective and acts as a potentiator. Like the endogenous HCN activator cAMP, NB5 increases the current by shifting the voltage dependency of the channel to the right but differently from cAMP which acts intracellularly and is not subtype-specific, its action is exerted from the extracellular side and is restricted to HCN4. NB5 exhibits additional properties which make it suitable for scientific and therapeutic applications. In addition to its high affinity (tens of nanomolar), that guarantees the persistence of effect after washing out, its purification from *E. coli* is easy and reproducible and it is stable in saline solution. Most importantly, NB5 recognizes the heterotetrametric channel composed of HCN4:HCN1 as well as the native f-channel of SAN myocytes. This finding is important in view of a therapeutic application, since the exact composition of native pacemaker f-channels in terms of stoichiometry of HCN subtypes is not exactly known.

We evaluated the effectiveness of NB5 in hiPSC-CMs expressing HCN4, which enables for spontaneous activity^18^. Our findings showed that NB5 incubation led to a significant increase in spontaneous beating rate, along with shortening of systolic and diastolic intervals, indicating gain-of-function of pacemaker mechanism. Additionally, NB5 had no impact on amplitude or heterogeneity of contraction, ruling out the occurrence of arrhythmic events within the cellular layer after NB5 application. These data clearly indicate that NB5 increases automaticity of hiPSC-CMs in the absence of adverse effects on contractility or rhythmicity. Overall, these data further validate the ability of NB5 to activate HCN4 and to induce gain-of-function in the pacemaker mechanism.

### Noncanonical electromechanical coupling in HCN channels revealed by NB5

One unexpected finding of this study is that NB5 changes the voltage dependency of HCN4 by binding in the pore region (S5-S6 linker). The evidence, emerging from mutagenesis analysis and cell electrophysiology, is that it binds to the S5-S6 loop that is part of the so-called pore turret. Two residues, D448 and N456, crucially contribute to the binding site and are indispensable for NB5 functional action.

The finding that NB5 modulates voltage dependence *via* the S5-S6 loop challenges present understanding of structure-function correlates in HCN channels and possibly unveils a noncanonical “electromechanical” coupling between the movement of the VSD and the pore, specifically the S5-S6 loop, so far unexplored.

HCN channels are activated by voltage and their VSD senses the electrical field and mechanically transmits it to the pore. In the canonical pathway, pore opening is initiated by the S4 movement and propagates *via* the S4-S5 linker to the pore domain. Non-canonical gating, on the other hand, refers to several hydrophilic and hydrophobic interactions between the VSD and the pore domain that stabilize the closed conformation of the pore and are gradually released during voltage gating. When the tight coupling between the two domains is lost either for mutagenesis^2,22^ and/or protein delipidation^7^, the channel becomes voltage-independent and the pore constitutively open. On the contrary, when the coupling is reinforced, such as for example in the presence of the anesthetic propofol^22^, then the closed state is stabilized and the channel requires more energy, i.e. a more negative voltage, to open. Non-canonical coupling paths have been so far identified in the cytosolic ends of the TM domains and in the hydrophobic interface between the pore and the VSD^23–25^. In this context, the finding that the turret can control voltage-dependent gating opens the possibility to control gating from the extracellular side.

Also relevant is the observation that the effects of NB5 and cAMP are additive, indicating that they act through two independent pathways. The interest in this finding relates to the potential use of NBs in the clinic because their binding will not alter the physiological control on heart rate by catecholamines acting *via* the cAMP signaling.

### Potential use of NB5 in clinics

HCN4 dysfunctions lead to a wide spectrum of cardiac conditions grouped under the term of sinus node dysfunction. This includes symptomatic bradycardia and/or more complex arrhythmia, atrial fibrillation, and AV block. It is well established that most of the HCN4-related cardiac diseases are caused by a decrease in current, due to altered biophysical properties and/or reduced number of channels at the plasma membrane^20^. Most of the mutations found in patients are indeed LOF^4,20,26^. The finding that NB5 acts independently of the voltage sensor and of cAMP, guarantees that it will rescue mutant phenotypes, even if the mutation alters the canonical activation pathway within the protein. We have shown this already in a proof-of-concept experiment performed on a mutation of a tachy-brady patient with altered response to cAMP^21^.

Symptomatic bradycardia due to a decrease in HCN4 current could be worsened upon ageing of the SAN, a condition that further reduces I_f_ current density^27^. Decline in SAN activity and basal heart rate is a widespread medical condition in the aging population typically leading to artificial pacemaker implantation^28^. In a recent study, the incidence of symptomatic bradycardia among older adults was found to be 6.2% in an urban emergency department^29^. In this case, the HCN4 channel is not mutated but the number of channels at the plasma membrane decreases, and the heart rate is reduced. Addition of NB5 can rescue the phenotype by activating the current of channels at the plasma membrane, compensating for their low number.

The results showing that NB5 increases the native I_*f*_ current by increasing action potential firing in sinoatrial node myocytes prove that this NB, or its variants, have a potential clinical application in age-related symptomatic sinus bradycardia. Therapeutic applications of nanobodies have some limitations related to short serum half-life and rapid renal clearance. Despite these weaknesses, their superior properties in terms of small size, low immunogenicity, high stability and water solubility allowed for several of them to enter clinical trials^30^.

### Conclusions

Although our goal was to select state-dependent NB targeting the open state of the HCN4 pore, we eventually found that NB5 targets a very short extracellular loop next to the pore mouth. Given the small size of the extracellular loops in HCN4, this result is *per se* remarkable and highlights the unique capability of *in vitro* selection that allowed counter-selection against not structurally related competitors but HCN4 itself. Adoption of non-animal-derived recombinant antibody technology not only guarantees versatility and reproducibility but also alleviates ethical concerns posed by animal-derived antibodies^31^.

## Acknowledgements

We thank Nadia Mekrane for helping with the human myocytes’ experiments. This project has received funding from the European Research Council (ERC) under the European Union’s Horizon 2020 research and innovation program ERC-2023-SyG (grant agreement n° 101118744) (to A.M.); Fondazione Telethon grant n° GGP20021 (to A.M.); Ministero della ricerca (MUR) Progetti di Rilevante Interesse Nazionale (PRIN) 2022 grant n° 2022EMA8FA (to A.M.) and *Fondation Leducq*, grant TNE 19CV03 (to A.M., D.D. and M.E.M.)

## Authors contribution

ASS, AS performed library screening; RC, AP performed patch experiments on HEK293 cells and LMGP on mouse SAN cells; PM and RP performed (rabbit) experiments; PM analyzed the (rabbit) results; AMG contributed to (rabbit) study design; HB performed experiments on human CMs, ACM analyzed results; RC, DD, MEM, GT, AS and AM designed the experiments with myocytes, analyze and interpret data; AM conceived the study and wrote the manuscript

## Material and Methods

### HCN constructs for protein purification

rbHCN4 construct used is the one described in Saponaro et al, (2021).

### HCN4 protein expression and purification

Freestyle HEK293F cell cultures (Thermo Fisher) were transiently transfected with pEGA: HCN4 (1 μg per ml) according to the procedure detailed in Saponaro et al., 2021^11^, Saponaro, Sharifzadeh, et al., 2021^34^ and Saponaro et al., 2024^7^. Purified His_6x_-eGFP-TEVsite-HCN4 protein, is kept in solution in a buffer containing 200 mM NaCl, 20 mM HEPES pH 7.0 and detergents (LMNG-CHS) at the concentration of 0.002% (w/v).

### Isolation of nanobody binders from yeast surface-display library

Isolation of nanobody (NB) binders was performed using a yeast surface-display library approach as previously described (McMahon et al., 2018^10^). A naive yeast library (5 × 10^9^ yeast) was incubated at 25 °C in galactose-containing tryptophan drop-out (Trp-) medium for 48h to induce nanobody expression. Induced cells were washed and resuspended in selection buffer (20 mM HEPES pH 7.5, 200 mM NaCl, 0.1% (w/v) BSA, 5 mM Maltose). The first round of MACS selection began with a preclear step which involved passing the NB-expressing yeast through an LD column (Miltenyi) to remove NBs interacting with anti-Alexa Fluor 647 magnetic microbeads (Miltenyi), anti-GFP Alexa Fluor 647 conjugated antibody (Thermofisher, 1:200), and purified His_6x_-eGFP (0.5 µM), the tag fused to the antigen (HCN4). Yeast was incubated with the above-described reagents for 1h at 4 °C before passing through the LD column. HCN4-binding NBs were enriched with two cycles of positive selection briefly summarized as follows: two steps of magnetic activated cell sorting (MACS), followed by one of fluorescence activated cell sorting (FACS). In all these steps cells were resuspended and washed in the above-described selection buffer supplemented with 0.002% (w/v) LMNG-CHS detergent mixture. The antigen used to the positive selection is the purified His_6x_-eGFP-HCN4 protein displaying the pore in the open conformation^11^. The second cycle of positive selection was intercalated by rounds of MACS-based depletion with HCN4 in the “closed” pore conformation through an LD column. The latter steps were performed to potentially enrich the library of binders to the HCN4 open pore, in search of potential pore blockers. For MACS, both positive (LS) and negative (LD) selection, the NB-expressing yeast (5 × 10^9^ in the first round and 3 × 10^7^ in the following rounds) was incubated, first, with 250 nM His_6x_-eGFP-HCN4 and anti-GFP-Alexa Fluor 647 conjugated antibody (1:200) for 1h at 4 °C, and then with anti-Alexa Fluor 647 magnetic microbeads for 1h at 4 °C. After each of the two incubations, a washing step was performed to ensure the removal of the labeling reagents not bound. Labeled yeast cells were passed through an either LS or LD column (Miltenyi), washed, and eluted by removing the magnetic field. The yeast cells eluted from LS column, or the ones washed from LD column were grown in glucose-containing Trp-medium for 24h at 30 °C to recover. Induction of NB expression was then repeated by incubation in galactose Trp-medium as described above. For FACS, the NB-expressing yeast (3 × 10^7^) was incubated with 250 nM His_6x_-eGFP-HCN4 for 1h at 4 °C. Removal of unbound antigen was ensured by a washing step performed immediately after incubation. Yeast cells binding to HCN4 via NB were then selected via flow cytometry following the green signal of the excited eGFP fused to HCN4. The sorted yeast cells were grown in glucose-containing Trp-medium for 24h at 30 °C to recover. Induction of NB expression was then repeated by incubation in galactose Trp-medium as described above. Finally, after a further cleaning step (LD MACS) to remove NB interacting with anti-Alexa Fluor 647 magnetic microbeads, anti-GFP Alexa Fluor 647 conjugated antibody, and purified His_6x_-eGFP, a final LS MACS was performed with HCN4 displaying the open pore as antigen. High affinity HCN4 NB binders were selected by plating yeast as single colonies for plasmid isolation and NB cDNA sequencing. Ten individual NBs were identified. Progressive enrichment of HCN4-selective NB binders was monitored via analytical FACS using 7×10^6^ NB-expressing cells pre-incubated 250 nM His_6x_-eGFP-HCN4 for 1h at 4 °C.

### Nanobody expression and purification

Isolated nanobody cDNAs were cloned into the periplasmic expression vector pET26b containing a C-terminal His_6x_ tag and were expressed in *Escherichia coli* Rosetta strain (EMD Millipore). Cells were grown at 37 °C in Terrific Broth medium (Research Products International) to an OD_600_ of 0.6 and induced with 1 mM isopropyl-1-thio-β-D-galactopyranoside (SIGMA) at 25 °C for 15h. Cells expressing NBs were collected by centrifugation and resuspended in lysis buffer (500 mM Sucrose, 200 mM Tris pH 8, 0.5 mM EDTA) and then osmotically shocked by diluting them three-times in a water solution containing 5 mM MgCl_2_ with 1h of stirring to release periplasmic NBs. The lysate was brought to a concentration of 100 mM NaCl and then cleared by centrifugation (14,000 g for 30 min at 4 °C). NBs were purified by affinity chromatography using HisTrap HP (GE Healthcare) pre-equilibrated with a high-salt buffer (500 mM NaCl, 20 mM HEPES pH 7.5-, and 20-mM Imidazole). The resin was washed with 10 CV of high-salt buffer, followed by 10 CV of a low-salt buffer (150 mM NaCl, 20 mM HEPES pH 7.5-, and 20-mM Imidazole). NBs were eluted in 150 mM NaCl, 20 mM HEPES pH 7.5-, and 300-mM Imidazole and loaded into a HiLoad 16/60 Superdex 75 prep grade size exclusion column (SEC) (GE Healthcare), pre-equilibrated with 150 mM NaCl, and 20 mM HEPES pH 7.5. All chromatographies were performed at 4 °C and monitored using the AKTApurifier UPC 10 fast protein liquid system (GE Healthcare).

### Electrophysiology on HEK293 cells

Patch clamp recordings were carried out on HEK293T cells or HEK293F cells. HEK293T cells were cultured in DMEM high glucose medium (Euroclone) supplemented with 10% FBS (Euroclone) and 1% Penicillin-Streptomycin (Sigma). HEK293F were grown in Freestyle medium (Thermo Fisher) supplemented with 10% FBS (Euroclone) and 1% Penicillin-Streptomycin (Sigma). Both cell types were grown at 37 °C degrees with 5% CO_2_. The cDNAs encoding full-length human HCN1, human HCN4, mouse HCN4 and hERG were previously cloned in pcDNA 3.1 mammalian expression vector, while full-length mouse HCN2 and rabbit HCN4 were previously cloned in pCI mammalian expression vector. When 70% confluent, cells were transfected (in a 35 mm petri dish) with Turbofect transfection reagent (Thermo Fisher). We used 1 µg of each construct and 0.3 µg of GFP-containing vector (pmax-GFP). For hHCN1 / rbHCN4 and rbHCN4 wt / rbHCN4 K531N heterotetramers, co-transfection of 1 µg of each cDNA was used. K531N, D448H and D448H_N456G mutations were introduced in rbHCN4 cDNA by site directed mutagenesis using the QuickChange II XL kit (Agilent Technologies). 24-48 h after transfection, cells were trypsinized and dispersed in 35 mm petri dishes. Single GFP^+^ cells were selected for patch clamp experiments that were carried out at room temperature. Currents were recorded in whole-cell configuration, using an ePatch (Elements srl) or a Dagan 3900A amplifier (Dagan Corporation). Signals acquired with the Dagan 3900A were digitized using a Digidata 1550B (Moelcular Devices). Patch-clamp signals were acquired with a sampling rate of 5 kHz and lowpass filter at 1 kHz. Patch pipettes were pulled from 1.5 mm O.D. and 0.86 mm I.D. borosilicate glass capillaries with a P-97 micropipette puller (Sutter, Novato, CA) and had resistances ranging from 3 to 6 MΩ. For all HCN channels recordings, patch pipettes were filled with a solution containing 10 mM NaCl, 130 mM KCl, 1 mM egtazic acid (EGTA), 0.5 mM MgCl_2_, 2 mM ATP (magnesium salt), and 5 mM HEPES–KOH buffer (pH 7.2), while the extracellular bath solution contained 110 mM NaCl, 30 mM KCl, 1.8 mM CaCl_2_, 0.5 mM MgCl_2_, and 5 mM HEPES–KOH buffer (pH 7.4). For hERG channel recordings patch pipettes were filled with a solution containing 130 mM KCl, 1 mM MgCl_2_, 5 mM egtazic acid (EGTA), 5 mM ATP (magnesium salt) and 10 mM HEPES-KOH buffer (pH 7.2), while the extracellular bath solution contained 140 mM NaCl, 4 mM KCl, 2.5 mM CaCl_2_, 10 mM Glucose, 5 mM HEPES-NaOH buffer (pH 7.45). Different volumes of nanobodies or, for all controls, the same buffer used to purify the nanobodies were added to either the pipette or bath solution to reach the concentration indicated for each experiment in the intracellular or extracellular solution, respectively. The NB purification buffer contains 150 mM NaCl, 20 mM HEPES and 10% (w/v) Glycerol at pH 7.5 supplemented with 1:1000 cOmplete^TM^, EDTA free (Sigma-Aldrich). Both buffer and nanobodies were stored at -80 °C until the day of the experiment where single-use aliquots were slowly thawed in ice. For washout experiments, NB5 was incubated in the petri dish for 2 minutes after which the NB-containing solution was removed, and patch clamp experiments were carried out during continuous perfusion of control solution for up to 70 minutes. Where indicated, adenosine 3′,5′-cyclic monophosphate (cAMP, Sigma-Aldrich) was added to the pipette solution from a previously prepared 100 mM stock solution (powder dissolved in milliQ water, pH adjusted to 7.0), to a final concentration of 30 µM. Single-use aliquots were prepared and stored at -20 °C until the day of the experiment.

To assess HCN channel activation curves, different voltage-clamp protocols were applied depending on the HCN subtype: for HCN1 holding potential was -20 mV (1 s), with steps from -30 mV to -120 mV (-10 mV increments, 3.5 s) and tail currents recorded at -40 mV (3.5 s); for HCN2, holding potential was -20 mV (1 s), with steps from -40 mV to -130 mV (-15 mV increments, 5 s) and tail currents recorded at -40 mV (5 s). For HCN4 and hHCN1 / rbHCN4 heterotetramers, holding potential was - 20 mV (1 s), with steps from -30 mV to -150 mV (-15 mV increments, 4.5 s) and tail currents recorded at -40 mV (4.5 s). For hERG holding potential was -80 mV (1s), with steps from -60 mV to +60 mV (Δ of +10 mV, 4 s) and tail currents were collected at -50 mV (6 s). Only cells in which a 1 GΩ seal or better was achieved were kept for analysis. For I/V plots currents were normalized to cell capacitance, indicated as I_SS_ (pA/pF). Neither series resistance compensation nor leak correction were applied.

Mean activation curves were obtained by fitting maximal tail current amplitude, plotted against the preconditioning voltage step, with the Boltzmann equation: y= 1/[1+exp((V−V_1/2_)/k)], where V is voltage, y the fractional activation, V_1/2_ the half-activation voltage, and k the inverse-slope factor in mV (k = -RT/zF). Mean activation curves were obtained by fitting individual curves from each cell to the Boltzmann equation and then averaging all the obtained values. The shift in V_1/2_ was plotted against NB5 concentration and fitted to a Hill equation (Y = Ymax*(1/(1 + (K_1/2_/x)nH))) to obtain K_1/2_ the concentration for half-maximal shift.

Activation and deactivation time constants (τ) were obtained by fitting a single exponential function: I=I_0_exp(-t/τ), to current traces obtained with the activation protocol described above. Deactivation time constants were obtained by fitting tail currents collected at -30, -45, -60 and -75 mV after a fully activating pulse at -135 mV (3.5 s).

To study NB5 state dependence, current traces were recorded at -90 mV using a repetitive activation protocol (1/30Hz)^16^ after the end of rundown (typically within 6min). NB-buffer or 2 µM NB5 were added to the bath solution while the cell was at its resting membrane potential (RPM, no voltage command). Currents were fitted with a single activation function as described above.

Data were analyzed with Clampfit (Molecular devices) and Origin (OriginLab) softwares and are presented as mean ± SEM. Statistical analysis were performed with the Student’s t-test for unpaired data, Student’s t-test for paired data or One-way ANOVA with Fisher’s test, as indicated for each experiment. Significance level was set to p = 0.05.

### Electrophysiology on mouse and rabbit sinoatrial node (SAN) cells

All experiments were carried out according to the ethical principles laid down by the French Ministry of Agriculture (agreement B91471109) and were performed conform to the guidelines from Directive 2010/63/EU of the European Parliament on the protection of animals. Experiments were performed on 2–3-month-old male New-Zeland rabbits (Charles River) kept under 12h:12h light/dark cycles with *ad-libitum* access to food and water. Rabbits were sedated with acepromazine maleate (Calmivet® 0.250 mg/kg intramuscularly) and sacrificed with Na^+^-penthobarbital (Euthasol® 70mg/kg intravenously + 1000U/kg heparin). The heart was quickly removed and washed in oxygenated Tyrode solution (in mmol/L: NaCl 130, KCl 5.4, NaH_2_PO_4_ 0.4, MgCl_2_ 0.5, CaCl_2_ 1.8, HEPES 25, glucose 22, pH adjusted to 7.4 with NaOH).

C57BL/6J mice aged 84-90 days from both genders were obtained from Charles River Laboratories, Italia S.r.l. Experimental procedures conformed to European and Italian laws (2010/63/EU D. and Lgs .2014/26) and were approved by the Animal Welfare Body of the University of Milan and by the Italian Ministry of Health (license n. 839C7.N.2BF). Animals were kept in pathogen-free conditions, with free access to food and water and were exposed to 12h light/dark cycles (light, 8 a.m. to 8 p.m.) in a thermostatically controlled room (21–22 °C).

### Single cell isolation

SAN pacemaker cells were isolated from C57BL6J mouse or New-Zeland rabbit hearts as previously described (DiFrancesco D. et al. 1986^35^). Mice were killed by cervical dislocation. Briefly, for both mice and rabbits, heart was excised and immersed in normal Tyrode solution (140 mM NaCl, 5.4 mM KCl, 1 mM MgCl_2_, 1.8 mM CaCl_2_, 5.5 mM D-glucose, and 5 mM HEPES (adjusted to pH 7.4 with NaOH) containing heparin pre-heated at 37 °C (mouse) or at room temperature (rabbit). The SAN tissue was excised by cutting along the crista terminalis and the interatrial septum and transferred into a low-Ca^2+^ solution containing 140 mM NaCl, 5.4 mM KCl, 0.5 mM MgCl_2_, 0.2 mM CaCl_2_, 1.2 mM KH_2_PO_4_, 50 mM taurine, 5.5 mM D-glucose, 1 mg/mL BSA, and 5 mM HEPES–NaOH (adjusted to pH 6.9 with NaOH). Enzymatic digestion was carried out for 25-30 min at 37 °C in the low-Ca^2+^ solution containing purified collagenase I and II (0.15 mg/mL Liberase medium Thermolysin, Roche, Mannheim, Germany) and elastase (0.5 mg/mL, Worthington, Lakewood, NJ, USA). The digested tissue was washed and transferred to a modified “Kraftbrühe” (KB) solution containing 100 mM K-glutamate, 10 mM K-aspartate, 25 mM KCl, 10 mM KH_2_PO_4_, 2 mM MgSO_4_, 20 mM taurine, 5 mM creatine, 0.5 mM EGTA, 20 mM D-glucose, 5 mM HEPES, and 1 mg/mL BSA (adjusted to pH 7.2 with KOH). Single cells were dissociated in Kraftbrühe solution at 37 °C by manual agitation using a flame-forged Pasteur’s pipette. To recover the automaticity of the SAN cells, Ca^2+^ was gradually reintroduced in the cell’s storage solution to a final concentration of 1.8 mM. Cells were used on the day of isolation.

### Patch-Clamp recordings of mouse and rabbit isolated SAN cells

Isolated mouse SAN cells were plated in glass bottom 35 mm Petri dishes (Greiner, bio-one) coated with 1-2 µg/mL Laminin (Merck Life Science S-r-l-, L2020). I_*f*_ current was recorded in whole-cell configuration using an ePatch amplifier (Elements, Cesena, Italy) at 34 °C and superfused with Tyrode solution supplemented with 2 mM BaCl_2_ and 2 mM MnCl_2_. Electrodes had a resistance of 3– 4 MΩ when filled with a solution containing 130 mM KCl, 10 mM NaCl, 0.5 mM MgCl_2_, 2 mM Mg-ATP, 1 mM EGTA and 5 mM HEPES (adjusted to pH 7.2 with KOH). To obtain I_*f*_ activation curve, a hyperpolarizing voltage step protocol was applied consisting of six steps from -135 mV (2.2 s) to -35 mV (9.2 s) with 20 mV and 2 s increments, starting from -35 mV holding potential and followed by a 1.2 s pulse at -135 mV.

Rabbit SAN cells were harvested in 35 mm Petri dishes, placed under the microscope and superfused at room temperature with Tyrode solution. Action potentials and I_*f*_ current were measured by perforated (escin 35 µM) and classical whole-cell patch clamp techniques and recorded using a Axopatch 200B patch clamp amplifier connected to Digidata 1550B interface (Molecular Devices). Electrodes had a resistance of 3-4 MΩ when filled with a solution containing 80 mM K-aspartate, 50 mM KCl, 1 mM MgCl_2_, 2 mM CaCl_2_, 5 mM EGTA, 5 mM HEPES, and 3 mM ATP-Na (adjusted to pH 7.2 with KOH). Pacemaker activity in isolated SAN cells was recorded before and after perfusion of 2 µM NB5. To measure I_*f*_ current and activation curve we used a protocol consisting of hyperpolarizing test pulses of 2 seconds between -35 and -135 mV in 10 mV increments from a holding potential of -35 mV followed by a 2 second step at -135 mV.

For both mouse and rabbit activation curve analysis, the current was obtained by subtracting the current at the beginning of the step to the steady state current. Mean activation curves were obtained by fitting maximal tail current amplitude, plotted against the preconditioning voltage step, with the Boltzmann equation, as described above.

### hiPSC maintenance and 2D cardiac differentiation

A previously characterized male healthy control hiPSC line (named RDES) was used for this study and maintained in cell culture as described in a previous publication^36^. Once the cells reached approximately 70–80% confluence, a second layer of reduced growth factor Matrigel (0.04 mg, Corning) was applied in StemFlex medium. Differentiation was carried out using a cardiac sheet protocol adapted for spontaneously beating ventricular-like hiPSC-CMs^36^. Starting on day 9, the medium was replaced every two days with RPMI 1640 + B27 supplemented with insulin, along with T3 hormone (3,3′,5-Triiodo-L-Thyronine) (Sigma, T2877) and liposoluble cAMP (N6,2′-O-Dibutyryladenosine 3′,5′-cyclic monophosphate sodium salt) (Sigma, D0627), both known to enhance hiPSC-CM electrophysiological and morphological features^37,38^.

### Contractility assay on human cardiac sheets

Paired contractility experiments on 2D spontaneously beating cardiac sheet were performed on day 37 in 6-well plates using a Zeiss LMS800 confocal microscope with a 10X lens in Tyrode’s solution at 37°C. 25 second edge capture videos were recorded of spontaneously contracting regions before and after incubation of 2µM NB5 for 2 min. Video data were processed using Zen Software (Zeiss) to extract raw recordings. A custom MATLAB script was then used to convert video frames into signals, which were subsequently analyzed to characterize contractility properties as previously reported^36,39,40^.

