## Supplementary Figures and Tables for "Animal-free recombinant nanobody rescues HCN4 channel deficit in sinus node dysfunction"

**Supplementary materials (Sharifzadeh et al)**

**Figures S1-S8**

**Table S1-S5**

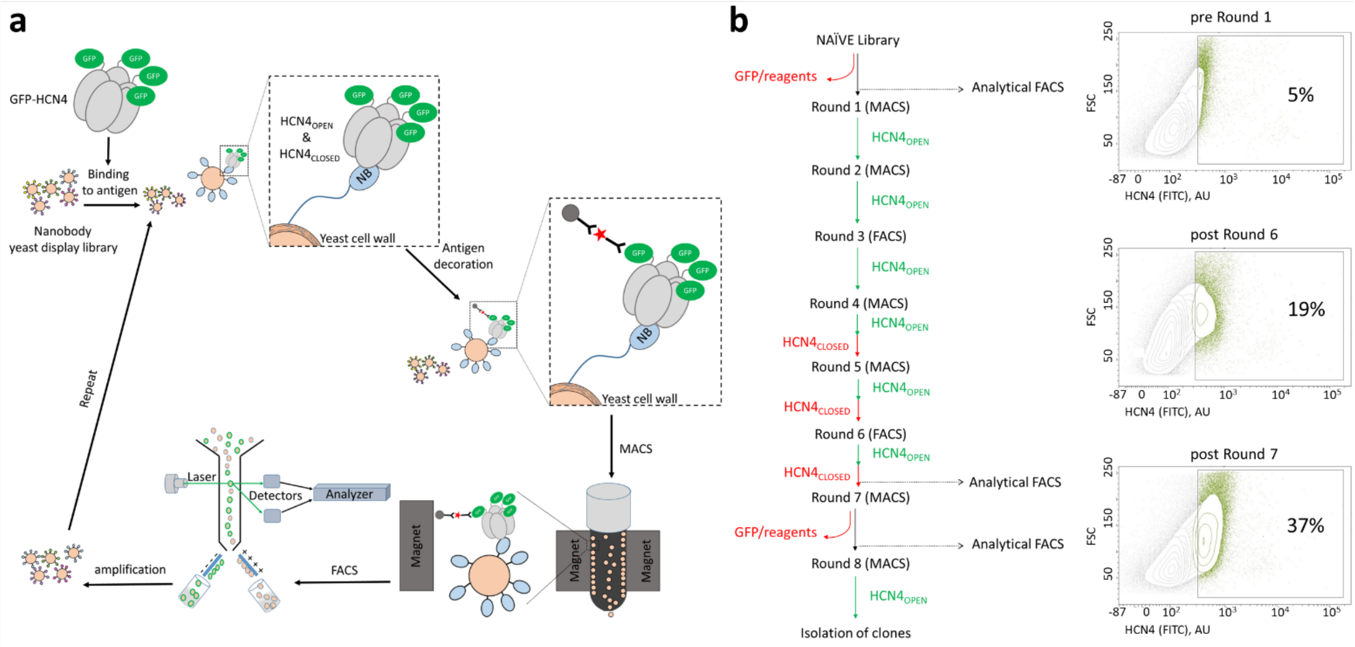

**Figure S1. Screening of a yeast-display synthetic nanobody library with purified HCN4 protein as antigen. a.** Schematic drawing of the screening process. Binding to antigen: the yeast-display nanobody library is exposed to purified GFP-HCN4. Inset: details of GFP-HCN4 interaction with a NB expressed on the surface of a yeast cell. $HCN4_{open}$ and $HCN4_{closed}$ refer to the antigen purified with the pore in the open and closed conformation, respectively. Antigen decoration included 1) anti-GFP antibody conjugated with Alexa 647 red star) and 2) anti-Alexa 647 antibody conjugated with a magnetic microbead (grey sphere). Inset: details of the double immunodecoration. Magnetic-Activated Cell Sorting (MACS) used to select NBs binding to HCN4. Fluorescence-Activated Cell Sorting (FACS): yeast cells interacting with the channel are sorted by means of the fluorescent signal of GFP-HCN4. **b.** Workflow of the selection procedure including two rounds of depletion of NBs binding to GFP and microbeads (GFP/reagents) and 8 rounds of MACS or FACS, alternating positive selection (using $HCN4_{OPEN}$ as antigen) and negative selection (using $HCN4_{CLOSED}$ as antigen). To monitor the enrichment of the library, three analytical FACS were performed, as indicated, before round 1, and after rounds 6 and 7. The side scatter (SSC) signal is plotted against the green fluorescence signal of HCN4 (509 nm) expressed in arbitrary units (AU). Green positive yeast cells are boxed and colored green. The first analytical FACS shows that 5% of the library interacts with $HCN4_{OPEN}$. After six rounds of positive selection, intercalated by three negative selections, the percentage of $HCN4_{OPEN}$ yeast binders increased by around four-fold (19%). After the last positive selection (round 7) this value reached 37%.

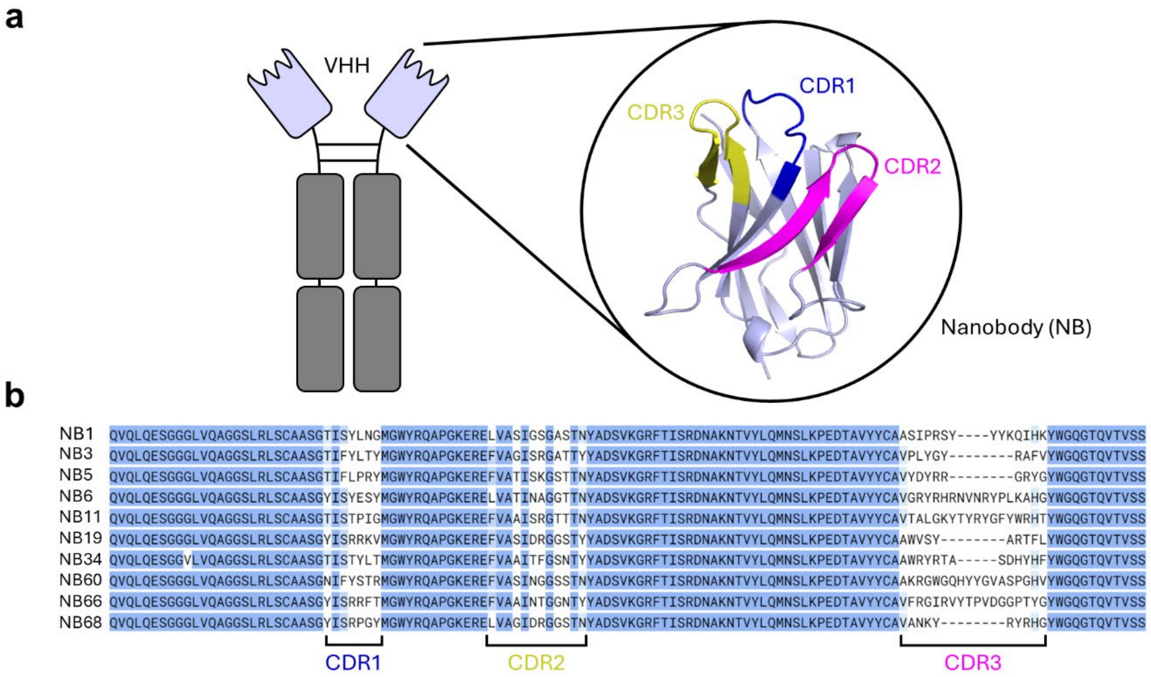

**Figure S2. Ten unique nanobodies isolated from the library. a.** Cartoon representation of a Camelid IgG heavy chain only antibody (HcAbs) each comprising two heavy chains (dark grey) that are linked by disulfide bonds and one variable domain (VHH, light blue). The isolated VHH region is referred to as "nanobody (NB)". Blow-up showing ribbon representation of NB5 obtained with AlphaFold2 Colab[32,33]. Complementarity-Determining Regions (CDR1, 2, 3) are highlighted in blue, magenta and yellow, respectively. **b.** Sequence alignment of the ten nanobodies isolated at the end of the selection process. Residues are highlighted with a blue color gradient by percentage identity (no color < 30% identity, light blue 60% = identity, blue = 70% identity, darkest blue = 100% identity), which measures the number of identical residues in relation to the length of the alignment. Framework regions are conserved while CDR1, 2, and 3 confer antigen-binding specificity.

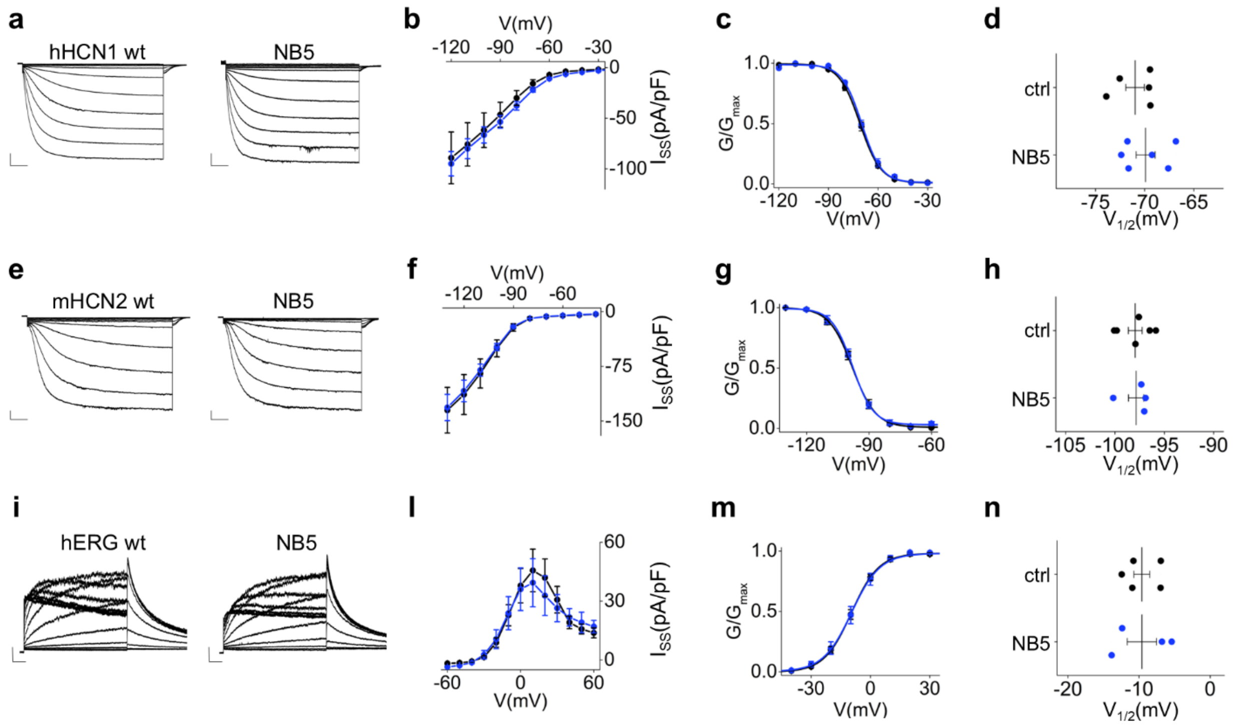

**Figure S3. NB5 has no effect on hHCN1, mHCN2 and hERG channels**. **a.** Representative whole-cell currents of hHCN1 wt recorded in control solution (left) or in presence of 20 µM NB5 in the extracellular solution (right). Traces shown are from -20 mV to -120 mV (see Material and Methods for voltage step protocol). Scale bars: 250 pA and 500 ms. **b.** I/V relationships of hHCN1 wt in control solution (black) or in presence of 20 µM NB5 in the extracellular solution (blue). Data are mean ± SEM. Current density values (in pA/pF), indicated as $I_{SS}$ (current at steady state), recorded at -120 mV, ctrl = -89.1 ± 25.6 (n = 4), +NB5 = -94.9 ± 12.0 (n = 6) are not statistically different ([§]p=0.8, Student's T-test). **c.** Activation curves obtained from hHCN1 wt currents in control solution (black circles) or in presence of 20 µM NB5 in the extracellular solution (blue circles). Data fit to the Boltzmann equation (see Material and Methods) are plotted as solid lines. Data points are mean ± SEM. Individual $V_{1/2}$ values are shown in d. **d.** Half-activation voltages ($V_{1/2}$) of control (black circles, n=5) and NB5-treated (blue circles, n=6) cells expressing hHCN1 wt. Mean $V_{1/2}$ ± SEM for ctrl = -70.9 ± 0.9 mV, +NB5 -69.9 ± 0.9 mV. The shift in half activation voltage ($\Delta V_{1/2}$) ± SEM = 1.0 ± 1.4 mV. **e.** Representative whole-cell currents of mHCN2 wt recorded in control solution (left) or in presence of 20 µM NB5 in the extracellular solution (right). Traces shown are from -40 mV to -130 mV (see Material and Methods for voltage step protocol). Scale bars: 250 pA and 500 ms. **f.** I/V relationships of mHCN2 wt in control solution (black) or in presence of 20 µM NB5 in the extracellular solution (blue). Data are mean ± SEM. Current density values (in pA/pF), indicated as $I_{SS}$ (current at steady state), recorded at -130 mV, ctrl = -135.2 ± 31.3 (n = 6), +NB5 = -131.5 ± 34.4 (n = 8) are not statistically different ([§]p=0.9, Student's T-test). **g.** Activation curves obtained from mHCN2 wt currents in control solution (black circles) or in presence of 20 μM NB5 in the extracellular solution (blue circles). Data fit to the Boltzmann equation (see Material and Methods) are plotted as solid lines. Data points are mean ± SEM. Individual $V_{1/2}$ values are shown in h. **h.** Half-activation voltages ($V_{1/2}$) of control (black circles, n=6) and NB5-treated (blue circles, n=4) cells expressing mHCN2 wt. Mean $V_{1/2}$ ± SEM for ctrl = -97.9 ± 0.7 mV, +NB5 -97.8 ± 0.8 mV. The shift in half activation voltage ($\Delta V_{1/2}$) ± SEM = 0.1 ± 1.1 mV. **i.** Representative whole-cell currents of hERG wt recorded in control solution (left) or in presence of 20 μM NB5 in the extracellular solution (right). Traces shown are from -60 mV to +60 mV (see Material and Methods for voltage step protocol). Scale bars: 250 pA and 500 ms. **l.** I/V relationships of hERG wt in control solution (black) or in presence of 20 μM NB5 in the extracellular solution (blue). Data are mean ± SEM. Current density values (in pA/pF), indicated as $I_{SS}$ (current at steady state), recorded at +10 mV, ctrl = 45.6 ± 10.8 (n = 5), +NB5 = 39.4 ± 12.3 (n = 4) are not statistically different ([§]p=0.7, Student's T-test). **m.** Activation curves obtained from hERG wt currents in control solution (black circles) or in presence of 20 μM NB5 in the extracellular solution (blue circles). Data fit to the Boltzmann equation (see Material and Methods) are plotted as solid lines. Data points are mean ± SEM. Individual $V_{1/2}$ values are shown in n. **n.** Half-activation voltages ($V_{1/2}$) of control (black circles, n=5) and NB5-treated (blue circles, n=4) cells expressing hERG wt. Mean $V_{1/2}$ ± SEM for ctrl = -9.6 ± 1.1 mV, +NB5 = -9.6 ± 2.0 mV. The shift in half activation voltage ($\Delta V_{1/2}$) ± SEM = 0.01 ± 2.2 mV.

$V_{1/2}$, $\Delta V_{1/2}$, inverse slope factors (k) and number of cells (n) for each experiment shown are reported in Supplementary Table S2 along with the details on statistical analysis.

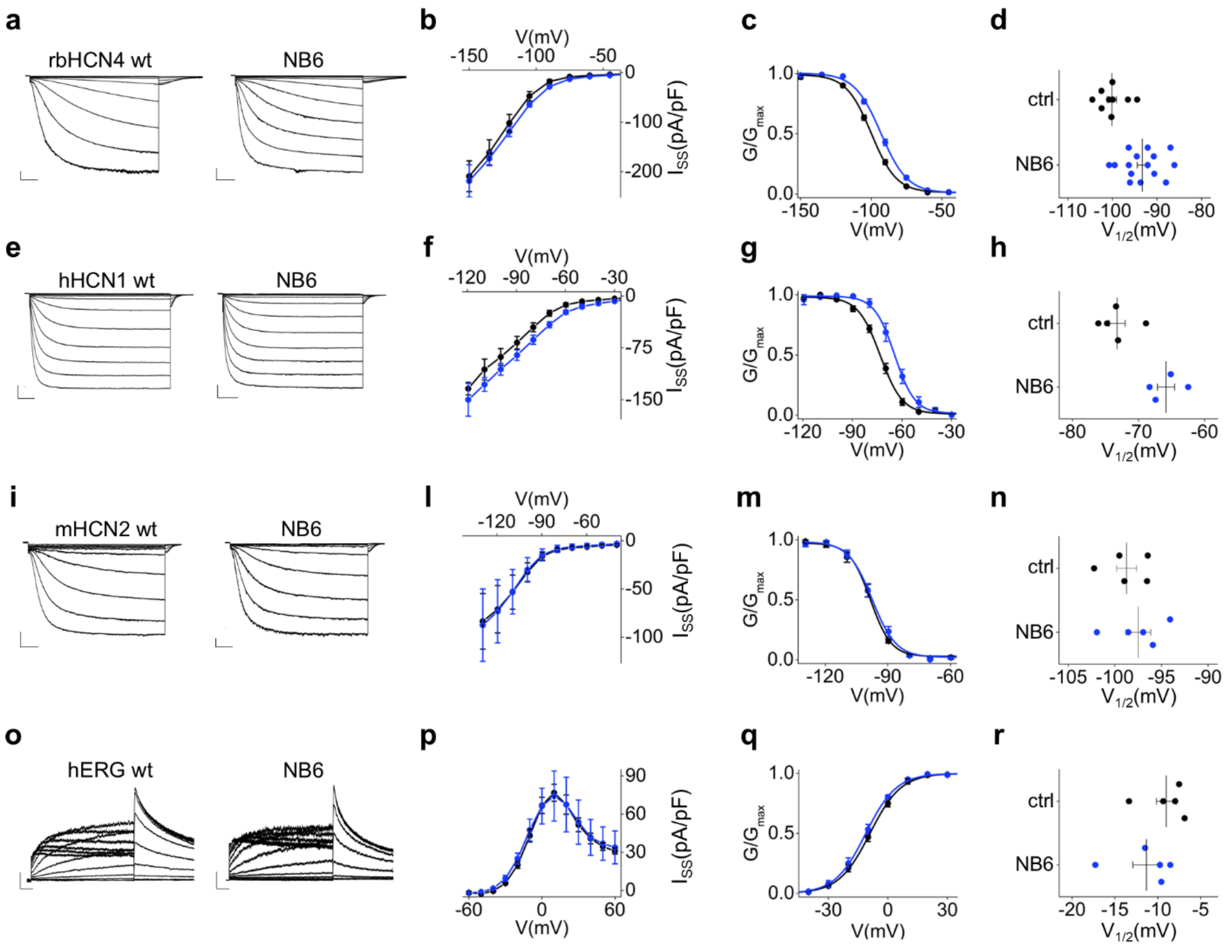

**Figure S4. NB6 acts on rbHCN4 and hHCN1, not on mHCN2 and hERG channels. a.** Representative currents of rbHCN4 in control (left) or in the presence of 20 μM NB6 in the extracellular solution (right). Traces shown are from -30 mV to -150 mV (see Material and Methods for voltage step protocol). Scale bars: 250 pA and 500 ms. **b.** I/V relationships reporting current density (in pA/pF) at steady state ($I_{SS}$), control (black), NB6 (blue). Data are mean ± SEM. Values at -150 mV: ctrl = -209.3 ± 31.0 (n = 6), +NB6 = -204.8 ± 30.0 (n = 11) are not statistically different ($^{§}$p=0.9, Student's T-test). **c.** Activation curves, control (black circles), NB6 (blue). Data fit to the Boltzmann equation (see Material and Methods) are plotted as solid lines. Data points are mean ± SEM. **d.** Half-activation voltages ($V_{1/2}$) of control (black, n=9) and NB6-treated (blue, n=15) cells. Mean $V_{1/2}$ ± SEM for ctrl = -100.2 ± 1.0 mV, +NB6 -93.3 ± 1.1 mV. Mean shift ($\Delta V_{1/2}$) ± SEM = 6.8 ± 1.6 mV. **e.** Representative currents of hHCN1 in control (left) or in the presence of 20 μM NB6 in the extracellular solution (right). Traces shown are from -20 mV to -120 mV. Scale bars: 250 pA and 500 ms. **f.** I/V relationships of hHCN1 in control (black) or with NB6 (blue). Data are mean ± SEM. Values at -120 mV, ctrl = -134.0 ± 9.0 (n = 5), +NB6 = -150.0 ± 23.8 (n = 5) are not statistically different ($^{§}p=0.5$, Student's T-test). **g.** Activation curves, control (black circles), NB6 (blue circles). Data fit to the Boltzmann equation are plotted as solid lines. Data points are mean ± SEM. **h.** $V_{1/2}$ of control (black, n=5) and NB6-treated (blue, n=4) cells expressing hHCN1. Mean $V_{1/2}$ ± SEM for ctrl = -73.2 ± 1.2 mV, +NB6 -64.9 ± 1.0 mV. Mean shift ($\Delta V_{1/2}$) ± SEM = 8.4 ± 1.7 mV. **i.** Representative currents of mHCN2 in control (left) or in the presence of 20 µM NB6 (right). Traces shown are from -40 mV to -130 mV. Scale bars: 250 pA and 500 ms. **l.** I/V relationships of mHCN2 in control (black) or with NB6 (blue). Data are mean ± SEM. Values at -130 mV, ctrl = -83.7 ± 28.8 (n = 4), +NB6 = -87.6 ± 37.5 (n = 4) are not statistically different ($^{§}p=0.9$, Student's T-test). **m.** Activation curves, control (black circles), NB6 (blue circles). Data fit to the Boltzmann equation are plotted as solid lines. Data points are mean ± SEM. **n.** $V_{1/2}$ of control (black, n=5) and NB6-treated (blue, n=5) cells expressing mHCN2. Mean $V_{1/2}$ ± SEM for ctrl = -98.7 ± 1.1 mV, +NB6 -97.5 ± 1.3 mV. Mean shift ($\Delta V_{1/2}$) ± SEM = 1.3 ± 1.7 mV. **o.** Representative currents of hERG in control (left) or with 20 µM NB6 (right). Traces shown are from -60 mV to +60 mV. Scale bars: 250 pA and 500 ms. **p.** I/V relationships of hERG in control (black) or with NB6 (blue). Data are mean ± SEM. Values at +10 mV, ctrl = 76.7 ± 6.7 (n = 5), +NB6 = 74.2 ± 19.6 (n = 4) are not statistically different ($^{§}p=0.9$, Student's T-test). **q.** Activation curves, control (black circles), NB6 (blue circles). Data fit to the Boltzmann equation are plotted as solid lines. Data points are mean ± SEM. **r.** $V_{1/2}$ of control (black, n=5) and NB6-treated (blue, n=5) cells expressing hERG. Mean $V_{1/2}$ ± SEM for ctrl = -9.0 ± 1.2 mV, +NB6 = -11.3 ± 1.6 mV. Mean shift ($\Delta V_{1/2}$) ± SEM = -2.3 ± 1.9 mV.

$V_{1/2}$, $\Delta V_{1/2}$, inverse slope factors (k) and number of cells (n) for each experiment shown are reported in Supplementary Tables S1 and S2 along with the details on statistical analysis.

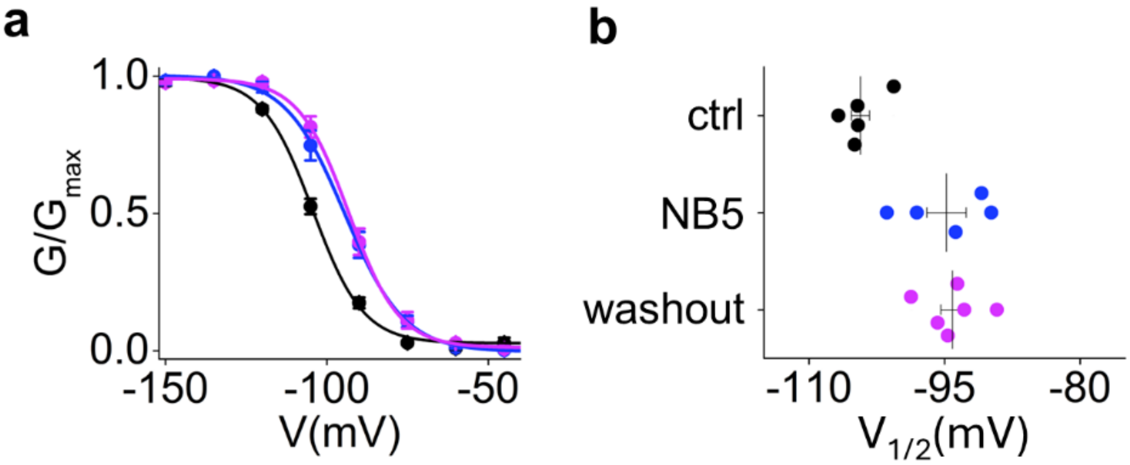

**Figure S5. NB5 effect on rbHCN4 is persistent after washout. a.** Activation curves obtained from rbHCN4 wt currents in control solution (black circles), in continuous presence of 2 μM NB5 in the extracellular solution (blue circles) or pre-incubated for 2 minutes with 2 μM NB5 in the extracellular solution and recorded during perfusion without NB (washout, magenta circles) (see Material and Methods). Data fit to the Boltzmann equation (see Material and Methods) are plotted as solid lines. Data points are mean ± SEM. Individual $V_{1/2}$ values are shown in **b.** Half-activation voltages ($V_{1/2}$) of control (black circles, n=5) or NB5-treated cells expressing rbHCN4 either continuously throughout the patch clamp session (blue circles, n=5) or during washout (magenta circles, n=6). Mean $V_{1/2}$ ± SEM for ctrl = -104.3 ± 1.0 mV, +NB5 = -94.8 ± 2.2 mV, washout = -94.1 ± 1.3 mV.

$V_{1/2}$, $\Delta V_{1/2}$, inverse slope factors (k) and number of cells (n) for each experiment shown are reported in Supplementary Table S3 along with the details on statistical analysis.

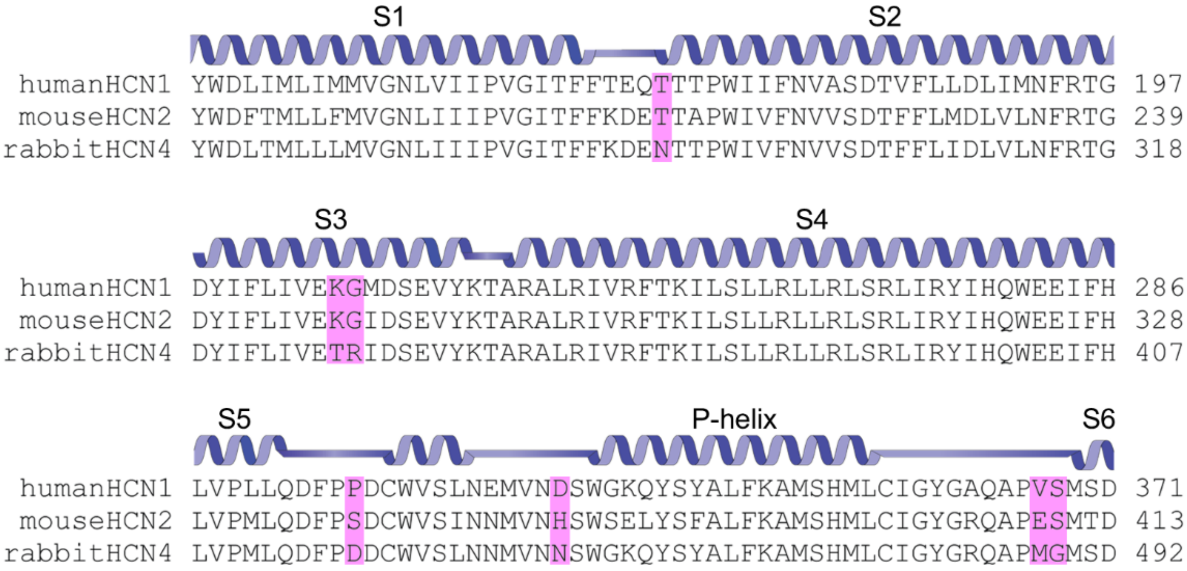

**Figure S6. rbHCN4 specific residues in extracellular regions.** Secondary structure elements and multiple sequence alignment of the extracellular portions and their flanking regions of human HCN1 (Gene ID: 348980), mouse HCN2 (Gene ID: 15166) and rabbit HCN4 (Gene ID: 100009452). Residues specific for rbHCN4 in the extracellular domains are highlighted in magenta.

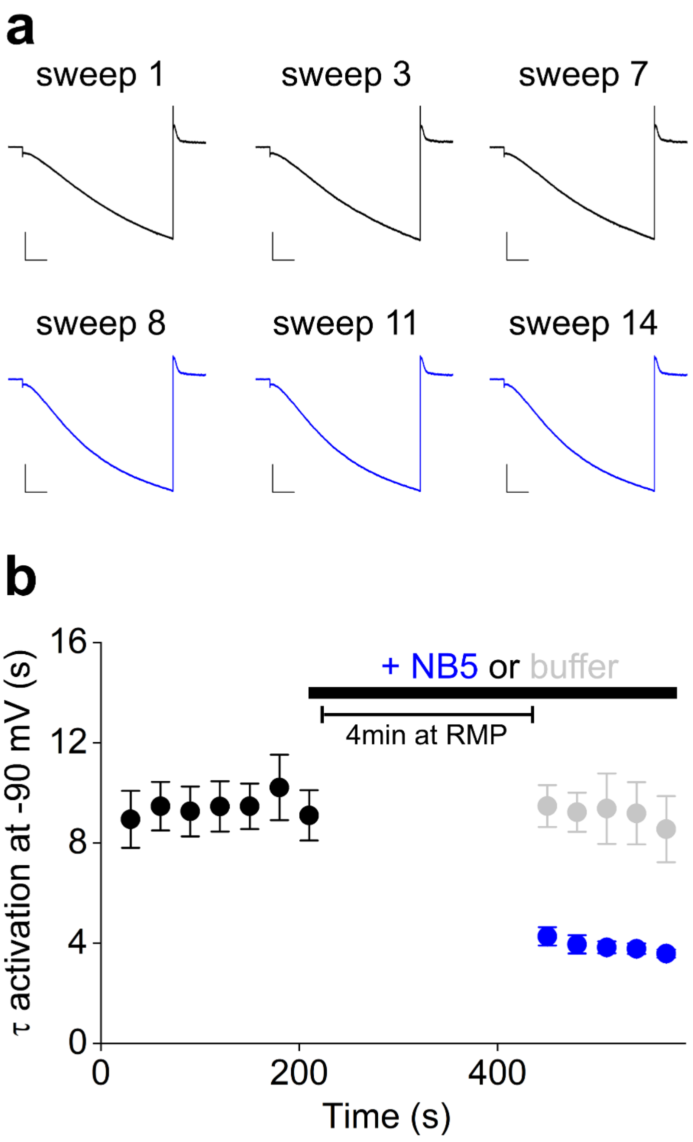

**Figure S7. The effect of NB5 is not state-dependent a.** Current traces of rbHCN4 at -90 mV recorded before (black) and after (blue) the exposure (4 minutes at RMP) of the cells to 2 μM NB5. **b.** Activation kinetics at -90 mV before (black) and after (blue) 4 minutes from the addition of 2 μM NB5 (blue) to the bath solution (black horizontal bar). Kinetics of control cells were measured in the same way using the NB buffer (grey) as treatment. Data points are mean ± SEM. During perfusion of either NB5 or buffer, any applied voltage command was removed leaving the cells at their resting membrane potential (RMP), usually ranging from -20 to -30 mV.

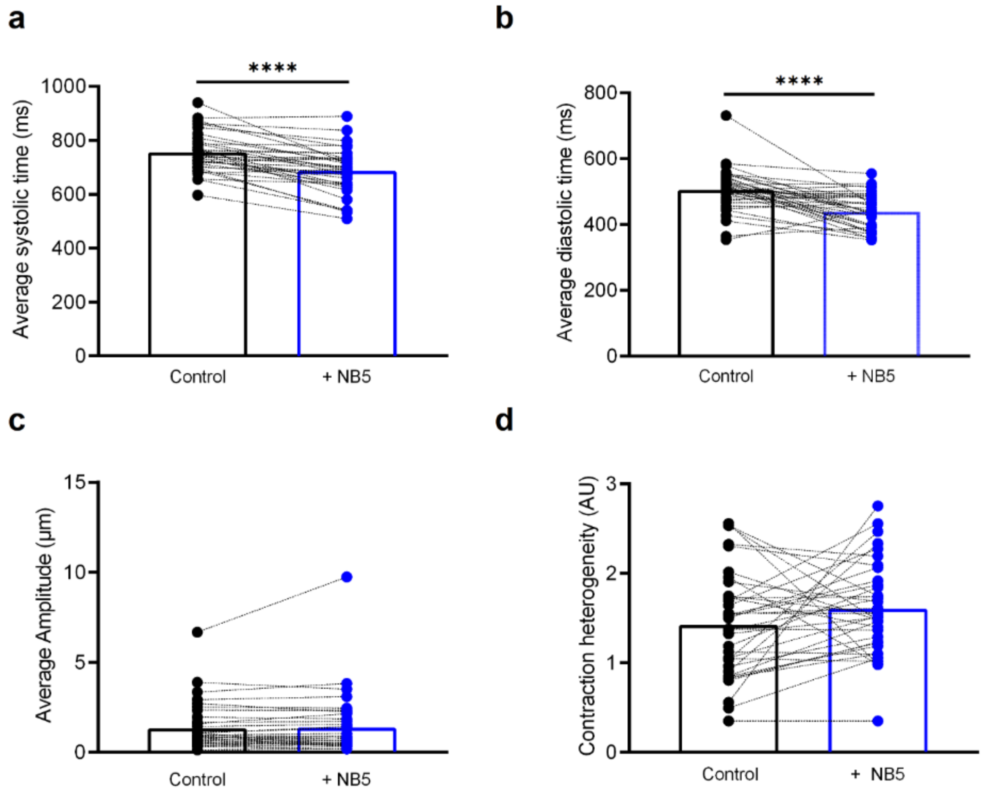

**Figure S8. NB5 causes shorter systole/diastole cycle in human cardiac sheet.**

**a-b.** Before-after graphs of the average systolic and diastolic times (ms) of spontaneously beating cardiac sheets before and after addition of 2 μM NB5. **c.** Before-after graph of the average amplitude (μm) of spontaneously beating cardiac sheets containing hiPSC-CMs before (ctrl) and after addition of 2 μM NB5 (NB5). **d.** Before-after graph of the contraction heterogeneity (AU) of spontaneously beating cardiac sheets containing hiPSC-CMs before and after addition of 2 μM NB5. Empty columns show mean values. Statistical analysis performed with Wilcoxon paired test (****$p < 0.0001$).

**Table S1**: **Functional effect of NBs on wt rbHCN4 in HEK293T cells.**

Comparison of two populations of cells, treated and not treated with NBs: extra and intra refer to the addition of the NB to the extracellular bath solution or to the pipette solution, respectively. All NBs were 20 μM (except for NB19 intra that was 10 μM, indicated by #). NB1 was unstable in extra and intra solutions and NB34 was unstable in intra solution. Half activation voltage ($V_{1/2}$) and inverse slope factor (k) values were obtained by fitting data to a Boltzmann function (see Material and Methods). n: number of cells; $\Delta V_{1/2}$: NB-induced shift in $V_{1/2}$, in mV. Each set of experiments contains data from control and NB measured on the same day. Values are reported as mean ± S.E.M., p values were calculated by Student's two-tailed unpaired T-test compared to wt rbHCN4 in control condition (without NB). P values highlighted in grey are not significant (> 0.05); N.D. not determined.

| | | **rbHCN4 wt** | | | **+20 μM NB** | | | |
|---|---|---|---|---|---|---|---|---|
| | | **$V_{1/2}$ (mV) ± SEM** | **k (mV) ±** | **n** | **$V_{1/2}$ (mV) ± SEM** | **k (mV) ±** | **n** | **$\Delta V_{1/2}$ (mV) ± SEM** |
| **extra** | **NB1** | N.D. | N.D. | | N.D | N.D. | | |
| | **NB3** | -103.7 ± 1.4 | 9.7 ± 0.6 | 7 | -95.2 ± 1.5 $^{p=0.013}$ | 8.9 ± 0.7 | 7 | 8.6 ± 2.0 |
| | **NB5** | -104.8 ± 0.8 | 9.9 ± 0.8 | 12 | -95.7 ± 0.9 $^{p=1.9E-7}$ | 9.9 ± 0.9 | 10 | 9.1 ± 1.2 |
| | **NB6** | -100.2 ± 1.0 | 9.1 ± 0.5 | 9 | -93.3 ± 1.1 $^{p=4.5E-4}$ | 9.4 ± 0.3 | 15 | 6.8 ± 1.6 |
| | **NB11** | -101.6 ± 0.6 | 9.4 ± 1.1 | 4 | -92.8 ± 1.0 $^{p=2.0E-4}$ | 8.6 ± 0.2 | 5 | 8.7 ± 1.2 |
| | **NB19** | -103.3 ± 1.2 | 10.4 ± 0.3 | 7 | -102.1 ± 1.8 $^{p=0.6}$ | 9.4 ± 0.6 | 6 | 1.2 ± 2.1 |
| | **NB34** | -106.8 ± 1.8 | 9.8 ± 0.8 | 3 | -103.0 ± 1.5 $^{p=0.2}$ | 10.8 ± 1.0 | 3 | 3.8 ± 2.3 |
| | **NB60** | -101.3 ± 1.0 | 9.9 ± 0.7 | 11 | -98.0 ± 0.6 $^{p=0.02}$ | 9.8 ± 0.7 | 8 | 3.3 ± 1.3 |
| | **NB66** | -105.4 ± 2.2 | 7.9 ± 0.8 | 5 | -91.9 ± 1.8 $^{p=6.9E-4}$ | 8.1 ± 0.7 | 8 | 13.5 ± 2.8 |
| | **NB68** | -97.2 ± 0.8 | 9.4 ± 0.5 | 11 | -95.2 ± 1.6 $^{p=0.2}$ | 9.5 ± 0.7 | 8 | 2.0 ± 1.6 |
| **intra** | **NB19** # | -103.0 ± 1.6 | 10.8 ± 0.5 | 3 | -105.6 ± 1.4 $^{p=0.3}$ | 9.7 ± 0.7 | 4 | -2.7 ± 2.1 |
| | **NB34** | N.D. | N.D. | | N.D. | N.D. | | |
| | **NB68** | -103.9 ± 1.3 | 9.1 ± 0.5 | 3 | -99.9 ± 1.2 $^{p=0.1}$ | 10.0 ± 0.2 | 3 | 4.0 ± 1.8 |

**Table S2: Fitting parameters of the activation curves of hHCN1 wt, mHCN2 wt and hERG wt channels treated with nanobodies (NBs).**

Half activation voltage ($V_{1/2}$) and inverse slope factor (k) values obtained by fitting data to a Boltzmann function (see Material and Methods) in absence or presence of NBs at 20 µM added to the extracellular recording solution; n: number of cells tested in each condition. $\Delta V_{1/2}$: NB-induced shift in $V_{1/2}$, in mV. Each set of experiments contains data from controls and NB measured on the same day. Values are reported as mean ± S.E.M. §p values were calculated by Student's two-tailed unpaired T-test compared to the control condition (channel without NB). P values highlighted in grey are not significant (> 0.05).

| | | **control** | | | **+ NB 20 µM** | | | |
|---|---|---|---|---|---|---|---|---|
| | | **$V_{1/2}$ (mV) ± SEM** | **k(mV)±SEM** | **n** | **$V_{1/2}$ (mV) ± SEM** | **k (mV) ± SEM** | **n** | **$\Delta V_{1/2}$(mV)±SEM** |
| **hHCN1** | **NB3** | -73.5 ± 1.0 | 6.6 ± 0.6 | 6 | -71.3 ± 1.4 p=0.2 | 6.1 ± 0.8 | 6 | 2.1 ± 1.7 |
| | **NB5** | -70.9 ± 0.9 | 6.2 ± 0.1 | 5 | -69.9 ± 0.9 p=0.4 | 6.1 ± 0.4 | 6 | 1.0 ± 1.4 |
| | **NB6** | -73.2 ± 1.2 | 6.7 ± 0.3 | 5 | -64.9 ± 1.0 p=0.001 | 6.3 ± 0.3 | 4 | 8.4 ± 1.7 |
| | **NB11** | -73.4 ± 1.7 | 6.6 ± 0.6 | 6 | -66.1 ± 1.0 p=0.003 | 6.3 ± 0.6 | 6 | 7.4 ± 1.9 |
| | **NB60** | -76.1 ± 0.9 | 7.0 ± 0.3 | 6 | -72.8 ± 0.5 p=0.02 | 6.3 ± 0.7 | 4 | 3.3 ± 1.0 |
| | **NB66** | -75.8 ± 0.9 | 6.9 ± 0.3 | 6 | -72.7 ± 0.7 p=0.03 | 5.5 ± 0.8 | 5 | 3.1 ± 1.2 |
| **mHCN2** | **NB5** | -97.9 ± 0.7 | 5.6 ± 0.6 | 6 | -97.8 ± 0.8 p=0.9 | 5.0 ± 0.4 | 4 | 0.1 ± 1.1 |
| | **NB6** | -98.7 ± 1.1 | 5.1 ± 0.6 | 5 | -97.5 ± 1.3 p=0.5 | 5.6 ± 0.4 | 5 | 1.3 ± 1.7 |
| **hERG** | **NB5** | -9.6 ± 1.1 | 6.8 ± 0.4 | 5 | -9.6 ± 2.0 p=1.0 | 7.0 ± 0.9 | 4 | 0.01 ± 2.2 |

| | | | | | | | | |
|---|---|---|---|---|---|---|---|---|
| | **NB6** | -9.0 ± 1.2 | 7.8 ± 0.4 | 5 | -11.3 ± 1.6 $^{p=0.3}$ | 7.6 ± 0.3 | 5 | -2.3 ± 1.9 |
| | **NB11** | -3.8 ± 0.9 | 6.6 ± 0.3 | 5 | -4.2 ± 0.1 $^{p=0.7}$ | 6.5 ± 0.2 | 4 | 0.3 ± 1.1 |
| | **NB60** | -10.8 ± 0.4 | 6.8 ± 0.1 | 6 | -10.9 ± 2.5 $^{p=1.0}$ | 6.8 ± 0.4 | 3 | 0.0 ± 1.7 |
| | **NB66** | -10.8 ± 0.4 | 6.8 ± 0.1 | 6 | -5.4 ± 1.1 $^{p=8.7E-4}$ | 6.9 ± 0.2 | 5 | 5.5 ± 1.1 |

**Table S3: Fitting parameters of the activation curves of HCN4 channels treated with NB5.**

Half activation voltage ($V_{1/2}$) and inverse slope factor (k) values obtained by fitting data to a Boltzmann function (see Material and Methods) in absence or presence of NB5; [NB5]: concentration of NB5 added to the extracellular recording solution, in micromolar (μM); n: number of cells tested in each condition. $\Delta V_{1/2}$: NB-induced shift in $V_{1/2}$, in mV. Each set of experiments contains data from controls and NB measured on the same day.

Values are reported as mean ± S.E.M. *p<0.05 values were calculated by One-way ANOVA with Fisher's test compared to wt rbHCN4 in control solution (without NB); §p values were calculated by Student's two-tailed unpaired T-test compared to control condition (channel without NB).

| Construct | $V_{1/2}$ (mV) ± SEM control | k (mV) ± SEM control | n | [NB5] (μM) | $V_{1/2}$ (mV) ± SEM with NB5 | k (mV) ± SEM with NB5 | n | $\Delta V_{1/2}$ (mV) ± SEM |
|---|---|---|---|---|---|---|---|---|
| **rbHCN4 wt** | -104.8 ± 0.8 | 9.9 ± 0.8 | 12 | **20** | -95.7 ± 0.9 $^{§p=1.9E-7}$ | 9.9 ± 0.9 $^{§p=0.9}$ | 10 | 9.1 ± 1.2 |
| | -104.1 ± 1.0 | 9.5 ± 0.6 | 8 | **2** | -94.5 ± 0.2 $^{§p=7.1E-5}$ | 8.5 ± 1.3 $^{§p=0.4}$ | 4 | 9.5 ± 1.5 |
| | -102.5 ± 0.8 | 9.7 ± 0.6 | 6 | **0.2** | -95.8 ± 1.1 $^{§p=0.001}$ | 9.8 ± 0.4 $^{§p=0.9}$ | 4 | 6.7 ± 1.3 |
| | -110.4 ± 0.2 | 10.2 ± 1.1 | 4 | **0.02** | -106.5 ± 1.0 $^{§p=0.008}$ | 10.4 ± 1.1 $^{§p=0.9}$ | 4 | 3.9 ± 1.0 |
| | -103.4 ± 1.3 | 10.0 ± 0.5 | 7 | **0.004** | -101.7 ± 0.6 $^{§p=0.47}$ | 9.6 ± 1.4 $^{§p=0.7}$ | 3 | 1.7 ± 2.0 |
| | -103.2 ± 1.3 | 9.9 ± 0.5 | 7 | **0.002** | -104.2 ± 0.7 $^{§p=0.6}$ | 9.0 ± 0.7 $^{§p=0.3}$ | 5 | 1.0 ± 1.7 |
| **hHCN4 wt** | -121.2 ± 1.4 | 8.8 ± 0.7 | 8 | **2** | -112.0 ± 2.1 $^{§p=0.004}$ | 9.0 ± 0.7 $^{§p=0.9}$ | 4 | 9.2 ± 2.0 |
| **mHCN4 wt** | -102.0 ± 1.3 | 8.4 ± 0.6 | 7 | **2** | -102.9 ± 1.5 $^{§p=0.6}$ | 9.6 ± 1.2 $^{§p=0.4}$ | 7 | 0.9 ± 2.0 |
| **rbHCN4 wt** | -101.8 ± 0.7 | 9.2 ± 0.3 | 20 | | | | | |
| **rbHCN4 D448H** | -100.6 ± 1.1 | 8.6 ± 0.4 $^{*p=0.3}$ | 10 | | | | | 1.2 ± 1.2 $^{§p=0.3}$ |
| | -96.9 ± 1.0 $^{*p=0.06}$ | 8.9 ± 0.3 | 12 | **20** | -87.0 ± 2.3 $^{§p=1.8E-5}$ | 8.2 ± 0.3 $^{§p=0.2}$ | 5 | 9.9 ± 2.1 |
| | -97.6 ± 2.3 $^{*p=0.06}$ | 9.8 ± 0.6 | 4 | **10** | -89.1 ± 1.9 $^{§p=0.008}$ | 9.1 ± 0.9 $^{§p=0.6}$ | 7 | 8.5 ± 2.2 |

| | | | | | | | |
|---|---|---|---|---|---|---|---|---|
| | -97.9 ± 1.0 $^{*p=0.06}$ | 8.8 ± 0.4 | 9 | **2** | -91.2 ± 1.0 $^{\S p=0.001}$ | 8.5 ± 0.4 $^{\S p=0.4}$ | 11 | 6.7 ± 1.6 |
| | -97.5 ± 2.3 $^{*p=0.06}$ | 8.9 ± 0.6 | 4 | **0.2** | -93.4 ± 1.0 $^{\S p=0.2}$ | 8.2 ± 0.5 $^{\S p=0.3}$ | 5 | 4.1 ± 2.1 |
| | -97.6 ± 2.3 $^{*p=0.06}$ | 9.8 ± 0.6 | 4 | **0.02** | -96.3 ± 1.6 $^{\S p=0.7}$ | 8.2 ± 0.2 $^{\S p=0.08}$ | 3 | 1.3 ± 3.5 |
| | -96.9 ± 1.0 $^{*p=0.06}$ | 8.9 ± 0.3 | 12 | **0.002** | -97.7 ± 1.1 $^{\S p=0.7}$ | 8.6 ± 0.7 $^{\S p=0.7}$ | 5 | -0.8 ± 2.0 |
| **rbHCN4 D448H_ N456G** | -101.6 ± 1.1 | 9.0 ± 0.5 $^{*p=0.7}$ | 11 | | | | | 0.2 ± 1.1 $^{\S p=0.8}$ |
| | -101.6 ± 1.1 | 9.0 ± 0.5 | 11 | **20** | -98.8 ± 2.9 $^{\S p=0.2}$ | 8.7 ± 0.4 $^{\S p=0.8}$ | 4 | 2.8 ± 2.3 |
| | -101.6 ± 1.1 | 9.0 ± 0.5 | 11 | **2** | -100.8 ± 1.2 $^{\S p=0.7}$ | 9.9 ± 1.1 $^{\S p=0.4}$ | 4 | 0.8 ± 2.3 |
| | -101.6 ± 1.1 | 9.0 ± 0.5 | 11 | **0.2** | -100.6 ± 1.6 $^{\S p=0.7}$ | 9.6 ± 1.1 $^{\S p=0.6}$ | 4 | 1.0 ± 2.3 |
| **hHCN1 wt** | -73.0 ± 0.9 | 6.6 ± 0.4 | 12 | | | | | |
| **rbHCN4 wt** | -104.0 ± 0.9 | 9.7 ± 0.6 | 12 | | | | | |
| **hHCN1 wt /rbHCN4 wt** | -84.8 ± 1.6 $^{*p=2.5E-12}$ | 8.6 ± 0.6 | 14 | **2** | -72.2 ± 2.0 $^{\S p=4.1E-5}$ | 8.8 ± 0.7 $^{\S p=0.8}$ | 12 | 12.7 ± 2.5 |
| **mouse SAN** | -84.8 ± 0.6 | 11.2 ± 1.6 | 4 | **4** | -84.1 ± 1.1 $^{\S p=0.6}$ | 10.2 ± 1.3 $^{\S p=0.6}$ | 4 | 0.7 ± 1.2 |
| **rabbit SAN** | -108.9 ± 2.9 | 12.5 ± 1.5 | 9 | **2** | -97.3 ± 4.5 $^{\S p=0.04}$ | 14.8 ± 0.8 $^{\S p=0.2}$ | 7 | 11.6 ± 5.1 |
| **rbHCN4 wt** | -101.6 ± 1.5 | 10.2 ± 0.7 | 10 | | | | | |
| **rbHCN4 K531N** | -115.0 ± 1.0 | 9.7 ± 0.6 $^{*p=0.6}$ | 10 | | | | | -13.3 ± 1.7 $^{\S p=4.8E-7}$ |
| **rbHCN4 wt /rbHCN4 K531N** | -112.7 ± 1.0 | 10.3 ± 0.3 $^{*p=0.9}$ | 11 | | | | | -11.1 ± 1.6 $^{\S p=3.4E-6}$ |

| | | | | | | | | |
|---|---|---|---|---|---|---|---|---|
| | -112.7 ± 1.0 | 10.3 ± 0.3 | 11 | **2** | -94.7 ± 0.7 §p=1.8E-9 | 9.5 ± 0.7 §p=0.2 | 6 | 18.0 ± 1.4 |
| | -112.7 ± 1.0 | 10.3 ± 0.3 | 11 | **0.2** | -101.9 ± 1.1 §p=4.9E-7 | 10.5 ± 0.7 §p=0.8 | 12 | 10.8 ± 1.5 |
| **rbHCN4 wt** | -104.3 ± 1.0 | 8.4 ± 0.2 | 5 | **2** | -94.8 ± 2.2 §p=0.001 | 8.9 ± 0.4 §p=0.5 | 6 | 9.5 ± 2.2 |
| **2 min incub. + washout** | -104.3 ± 1.1 | 8.4 ± 0.2 | 5 | **2** | -94.1 ± 1.3 §p=4.0E-4 | 8.8 ± 0.8 §p=0.6 | 6 | 10.2 ± 2.2 |

**Table S4**: **Fitting parameters of the activation curves of rbHCN4 wt channel treated with NB5 and cAMP.**

Half activation voltage ($V_{1/2}$) and inverse slope factor (k) values obtained by fitting data to a Boltzmann function (see Material and Methods) in absence or presence of NB5 2 μM and/or cAMP 30 μM; n: number of cells tested in each condition. $\Delta V_{1/2}$ (shift from wt): shift in $V_{1/2}$ due to NB5, cAMP, or both from rbHCN4 wt in control condition (without NB and/or cAMP). $\Delta V_{1/2}$ (due to NB5): NB5-induced shift in $V_{1/2}$. Each set of experiments contains data from controls and NB measured on the same day.

Values are reported as mean ± S.E.M. *p<0.05 values were calculated by One-way ANOVA with Fisher's test compared to wt rbHCN4 in control solution (without NB). P values highlighted in red indicate statistical significance.

| channel | treatment | $V_{1/2}$ (mV) ± SEM | k (mV) ± SEM | n | $\Delta V_{1/2}$ (mV) ± SEM shift from wt | $\Delta V_{1/2}$ (mV) ± SEM due to NB5 |
|---|---|---|---|---|---|---|
| **rbHCN4 wt** | **control** | -104.7 ± 1.0 | 9.4 ± 0.6 | 9 | | |
| | **+2μM NB5** | -96.2 ± 1.6 $^{*p=7.4E-4}$ | 11.5 ± 0.7 $^{*p=0.07}$ | 6 | 8.5 ± 2.1 | 8.5 ± 2.1 |
| | **+30μM cAMP** | -86.8 ± 2.2 $^{*p=9.0E-8}$ | 10.6 ± 1.0 $^{*p=0.2}$ | 5 | 18.0 ± 2.3 | |
| | **+2 M NB5 +30 μM cAMP** | -76.2 ± 2.1 $^{*p=7.0E-12}$ | 10.4 ± 0.4 $^{*p=0.3}$ | 6 | 28.5 ± 2.2 | 10.6 ± 2.5 |

**Table S5**: **Activation and deactivation time constants ($\tau$) of channels treated with NB5 and/or cAMP.**

Mean activation and deactivation time constants ($\tau$) of rbHCN4 wt channel treated with 2 µM NB5 and 30 µM cAMP, hHCN1 / rbHCN4 heterotetramers treated with 2 µM NB5 and rbHCN4 wt / rbHCN4 K531N treated with 0.2 µM NB5. Time constants were calculated, at the indicated voltages (V), by fitting a single exponential function (see Material and Methods) to current traces obtained, with the activation and deactivation protocols described in the Material and Methods section, in absence or presence of NB5 and/or cAMP. n: number of cells tested in each condition; [NB5]: concentration of NB5 added to the extracellular recording solution, in micromolar (µM). N.D. not determined. Each set of experiments contains data from controls and NB measured on the same day.

Values are reported as mean ± S.E.M. *p<0.05 values were calculated by One-way ANOVA with Fisher's test compared to wt rbHCN4 in control solution (without NB). §p values were calculated by Student's two-tailed unpaired T-test compared to control condition (channel without NB).

| **Construct** | | **V (mV)** | **$\tau$ (s) ± SEM ctrl** | **n** | **[NB5] (µM)** | **$\tau$ (s) ± SEM + NB5** | **n** | **$\tau$ (s) ± SEM + cAMP** | **n** | **$\tau$ (s) ± SEM + NB5 + cAMP** | **n** |
|---|---|---|---|---|---|---|---|---|---|---|---|
| **rbHCN4 wt** | **activation** | -150 | 0.56 ± 0.04 | 6 | **2** | 0.79 ± 0.12 *p=0.06 | 6 | 0.61 ± 0.01 *p=0.7 | 3 | 0.46 ± 0.08 *p=0.4 | 3 |
| | | -135 | 1.1 ± 0.09 | 8 | **2** | 0.90 ± 0.05 *p=0.04 | 5 | 0.70 ± 0.02 *p=1.0E-4 | 7 | 0.63 ± 0.03 *p=4.5E-5 | 5 |
| | | -120 | 3.2 ± 0.63 | 8 | **2** | 1.5 ± 0.11 *p=0.007 | 5 | 1.0 ± 0.04 *p=4.6E-4 | 7 | 0.85 ± 0.02 *p=5.8E-4 | 5 |
| | | -105 | 7.1 ± 0.65 | 3 | **2** | 4.1 ± 0.72 *p=0.002 | 6 | 2.0 ± 0.23 *p=1.8E-5 | 5 | 1.3 ± 0.08 *p=4.8E-6 | 5 |
| | **deactivation** | -75 | 3.5 ± 0.56 | 8 | **2** | 4.3 ± 0.66 *p=0.3 | 5 | 5.2 ± 0.40 *p=0.07 | 4 | N.D. | |
| | | -60 | 2.1 ± 0.23 | 9 | **2** | 2.9 ± 0.41 *p=0.1 | 9 | 3.4 ± 0.45 *p=0.03 | 6 | N.D. | |
| | | -45 | 1.0 ± 0.11 | 10 | **2** | 1.2 ± 0.12 *p=0.3 | 11 | 1.9 ± 0.20 *p=1.4E-4 | 10 | 2.3 ± 0.15 *p=3.2E-6 | 6 |
| | | -30 | 0.43 ± 0.03 | 9 | **2** | 0.53 ± 0.03 *p=0.5 | 11 | 0.86 ± 0.14 *p=0.002 | 11 | 1.3 ± 0.13 *p=2.5E-6 | 6 |

| | | | | | | | |
|---|---|---|---|---|---|---|---|
| **hHCN1 wt /rbHCN4 wt** | activation | -150 | 0.23 ± 0.03 | 3 | **2** | 0.18 ± 0.03 $^{\S p=0.3}$ | 4 |
| | | -135 | 0.33 ± 0.04 | 11 | **2** | 0.19 ± 0.02 $^{\S p=0.01}$ | 8 |
| | | -120 | 0.43 ± 0.06 | 11 | **2** | 0.24 ± 0.03 $^{\S p=0.01}$ | 8 |
| | | -105 | 0.62 ± 0.09 | 11 | **2** | 0.30 ± 0.04 $^{\S p=0.01}$ | 9 |
| | deactivation | -40 | 0.38 ± 0.06 | 12 | **2** | 0.37 ± 0.05 $^{\S p=0.9}$ | 11 |
| **rbHCN4 wt** | activation | -150 | 0.51 ± 0.08 | 3 | | | |
| | | -135 | 0.92 ± 0.06 | 8 | | | |
| | | -120 | 2.0 ± 0.26 | 8 | | | |
| | deactivation | -40 | 0.56 ± 0.05 | 10 | | | |
| **rbHCN4 wt /rbHCN4 K531N** | activation | -150 | 0.79 ± 0.05 $^{*p=0.004}$ | 7 | **0.2** | 0.64 ± 0.03 $^{*p=0.2}$ | 5 |
| | | -135 | 1.5 ± 0.09 $^{*p=7.6E-5}$ | 10 | **0.2** | 1.0 ± 0.08 $^{*p=0.3}$ | 10 |
| | | -120 | 5.1 ± 0.84 $^{*p=5.7E-4}$ | 10 | **0.2** | 2.0 ± 0.20 $^{*p=0.9}$ | 11 |
| | deactivation | -40 | 0.75 ± 0.09 $^{*p=0.1}$ | 11 | **0.2** | 0.78 ± 0.08 $^{*p=0.07}$ | 10 |
